# Phylogenomic analysis reveals the basis of adaptation of *Pseudorhizobium* species to extreme environments

**DOI:** 10.1101/690347

**Authors:** Florent Lassalle, Seyed M. M. Dastgheib, Fang-Jie Zhao, Jun Zhang, Susanne Verbarg, Anja Frühling, Henner Brinkmann, Thomas H. Osborne, Johannes Sikorski, Francois Balloux, Xavier Didelot, Joanne M. Santini, Jörn Petersen

**Affiliations:** Department of Infectious Disease Epidemiology, Imperial College London, UK; MRC Centre for Global Infectious Disease Analysis, Imperial College London, London, UK; Research Institute of Petroleum Industry, Tehran, Iran; State Key Laboratory of Crop Genetics and Germplasm Enhancement, College of Resources and Environmental Sciences, Nanjing Agricultural University, Nanjing, China; Leibniz Institut DSMZ, Braunschweig, Germany; Institute for Structural & Molecular Biology, University College London, London, UK; UCL Genetics Institute, University College London, London, UK; School of Life Sciences, University of Warwick, Coventry, UK

**Author notes:** University of Bedfordshire, Luton, UK.

**Keywords:** *Rhizobium* sp. NT-26, genome taxonomy, clade-specific genes, ecological specialization, phylogenomics, pangenome analysis

## Abstract

The family *Rhizobiaceae* includes many genera of soil bacteria, often isolated for their association with plants. Herein, we investigate the genomic diversity of a group of *Rhizobium* species and unclassified strains isolated from atypical environments, including seawater, rock matrix or polluted soil. Based on whole-genome similarity and core genome phylogeny, we show that this group corresponds to the genus *Pseudorhizobium.* We thus reclassify *Rhizobium halotolerans, R. marinum, R. flavum* and *R. endolithicum* as *P. halotolerans* comb. nov., *P. marinum* comb. nov.*, P. flavum* comb. nov. and *R. endolithicum* comb. nov., respectively, and show that *P. pelagicum* is a synonym of *P. marinum*. We also delineate a new chemolithoautotroph species, *P. banfieldiae* sp. nov., whose type strain is NT-26^T^ (= DSM 106348^T^ = CFBP 8663^T^). This genome-based classification was supported by a chemotaxonomic comparison, with gradual taxonomic resolution provided by fatty acid, protein and metabolic profiles. In addition, we used a phylogenetic approach to infer scenarios of duplication, horizontal transfer and loss for all genes in the *Pseudorhizobium* pangenome. We thus identify the key functions associated with the diversification of each species and higher clades, shedding light on the mechanisms of adaptation to their respective ecological niches. Respiratory proteins acquired at the origin of *Pseudorhizobium* are combined with clade-specific genes to encode different strategies for detoxification and nutrition in harsh, nutrient-poor environments. Finally, we predict diagnostic phenotypes for the distinction of *P. banfieldiae* from other *Pseudorhizobium* species, including autotrophy and sensitivity to the azo dye Congo Red, which we experimentally validated.

## Introduction

Bacteria of the family *Rhizobiaceae* (*Alphaproteobacteria*) are usually soil-borne and found in association with plant roots, where they mostly rely on a saprophytic lifestyle degrading soil organic compounds and plant exudates, including aromatic compounds [8,9,48]. This particular versatility in using various organic compounds likely stems from the presence of some of the largest known sets of carbohydrate transporter genes in *Rhizobiaceae* genomes [38,73]. Some members of this taxon sometimes engage in a symbiotic or pathogenic relationship with a specific host plant, with the ability to switch to these lifestyles being determined by the presence of accessory megaplasmids in the bacterium [10,21,73]. It is more unusual to isolate them from an arsenic-containing rock in a sub-surface environment mostly devoid of organic matter such as the Granites goldmine in the Northern Territory, Australia [51]. Rock samples from this mine containing arsenopyrite (AsFeS) were used to enrich for and isolate organisms (designated NT-25 and NT-26) capable of using arsenite (oxidation state +3, i.e. As(III)) as the electron donor coupling its oxidation to arsenate (As(V)) with oxygen and using carbon dioxide as the sole carbon source [51]. 16S rRNA gene sequence analysis revealed that these strains were very closely related and likely belonging to the same species in the family *Rhizobiaceae*; they were provisionally named *Rhizobium* sp. [51]. However, recent advances in multi-locus sequence analysis (MLSA) and genome sequencing led to the recognition of the polyphyly of the genus *Rhizobium*. Subsequently, the taxonomy of *Rhizobiaceae* was largely revised, with many *Rhizobium* species being reclassified in newly created genera, including *Neorhizobium*, *Pararhizobium*, *Allorhizobium* and *Pseudorhizobium* [26,40–42] suggesting that the taxonomic status of NT-25 and NT-26 should be re-examined.

The strains NT-25 and NT-26 can withstand very high levels of arsenic (greater than 20 mM arsenite and 0.5 M arsenate), thanks to functions encoded notably by the arsenic-resistance (*ars*) genes [49]. In addition, they can gain energy from the oxidation of arsenite, with the mechanism in NT-26 studied in detail [5,6,16,49,50,52,70]. Analysis of the NT-26 genome sequence revealed other key genes involved in the resistance to arsenic, including the *phn*/*pst* genes, which encode phosphate-specific transporters with high affinity for phosphate but not for its structural analogue arsenate. This likely explains the high tolerance of NT-26 to arsenate [1]. The key determinants of arsenic metabolism and resistance, including the *aio*, *ars* and *phn/pst* genes, were primarily located on an accessory 322-kb megaplasmid distantly related to symbiotic plasmids of rhizobia. In addition, comparative genomics showed that the *aio* operon formed a stable genetic unit that can be found sporadically in a diverse set of bacteria – the closest relatives to these accessory genes residing in genomes of other members of the *Rhizobiaceae* [1,2].

Having identified the genetic features allowing NT-25 and NT-26 to live in harsh environments, we sought to investigate if this combination of adaptive determinants were only present in these ecologically specialized strains, or if they were the trademark of a wider taxon. We thus searched organisms closely related to NT-25/NT-26 based on their 16S rRNA sequence. Their closest relative, strain TCK [9], was selected for its ability to oxidize sulphur compounds, including hydrogen sulphide, sulphite and thiosulphate, a phenotype shared by NT-25 and NT-26 [1]. Other close relatives were strikingly all isolated from polluted environments: *R.* sp. strain Khangiran2 from a soil contaminated with petroleum, *R.* sp. strain Q54 from an arsenic-contaminated paddy soil; *R. flavum* strain YW14 from organophosphorus insecticide-contaminated soil [13] and *R. halotolerans* AB21 from soil contaminated with the detergent chloroethylene [10]. Interestingly, several pathways mediating resistance to these toxins or relating to their metabolism rely on the cellular respiration machinery, including oxidative degradation of noxious organic compounds, or the oxidation of arsenite. The more distantly related species *R. endolithicum* has the peculiar ability to live within a rock matrix [36], whereas the even more distant relatives *Pseudorhizobium pelagicum* and *R. marinum* live in open sea waters, a quite unusual feature within the *Rhizobiaceae* [18,29]. The environments these organisms were isolated from suggest bacteria in this group have a special ability to live in habitats that are chemically harsh and depleted in organic nutrients.

We sequenced the genomes of all known bacterial isolates closely related to *Rhizobium* sp. NT-26, resulting in eight complete or almost complete genome sequences. We complemented our dataset with all currently available complete or near-complete genome data for the bacterial families *Rhizobiaceae* and (closest relative) *Aurantimonadaceae*. We used this extensive dataset to compute a robust phylogenomic tree covering a broad taxonomic scope, which led us to the delineation of a new species, *Pseudorhizobium banfieldiae* sp. nov., and the reclassification of four species into the genus *Pseudorhizobium*. Using a phylogenetic framework, we analysed the distribution and history of all pangenome genes in this genus, and revealed key genetic innovations along its diversification history. Crucially, a large repertoire of respiratory chain proteins was acquired by the ancestor of *Pseudorhizobium* and later expanded in descendant lineages. The diversification of *Pseudorhizobium* species was then marked by their respective acquisition of unique metabolic pathways, providing each species with some specific detoxification mechanisms. Finally, we predicted and experimentally tested phenotypic traits that characterize and distinguish the studied species, providing at least one new diagnostic phenotype (sensitivity of *P. banfieldiae* to the azo dye Congo Red).

## Methods

### DNA extraction, genome sequencing and genome assembly

Two independent projects were conducted at University College London (UCL; London, UK) in collaboration with the Earlham Institute (Norwich, UK) and at the Leibniz Institute-DSMZ (Braunschweig, Germany) for the genome sequencing of strains *Rhizobium* sp. NT-25 (= DSM 106347) [51], *R. flavum* YW14^T^ (= DSM 102134^T^ = CCTCC AB2013042^T^ = KACC 17222^T^) [13], *R.* sp. Q54 (= DSM 106353), *R.* sp. TCK (= DSM 13828) [9] and Khangiran2 (= DSM 106339 = IBRC-M 11174). In addition, strains *R. halotolerans* AB21^T^ (= DSM 105041^T^ = KEMC 224-056^T^ = JCM 17536^T^) [10], *R. endolithicum* JC140^T^ (= DSM 104972^T^ = KCTC32077^T^ = CCUG64352^T^ = MTCC11723^T^ = HAMBI 2447^T^) [36] and *R.* sp. P007 [12] were sequenced only at the DSMZ.

For long-read sequencing, cells were cultured at UCL until stationary phase in a minimal salts medium (MSM) containing 0.08% yeast extract (YE) at 28°C [51]. Genomic DNA was extracted using the Wizard DNA Purification kit (Promega, Madison, Wisconsin) according to the manufacturer’s instructions. Quality of the genomic DNA was assessed as described in the Supplementary Methods. DNA libraries were prepared for sequencing at the Earlham Institute on the PacBio RSII platform using C4-P6 chemistry with one SMRT cell per genome. This generated 82–174 ×10^3^ long reads per genome (mean: 139 ×10^3^), representing 0.332–1.26 ×10^9^ bp (mean: 0.982 ×10^9^). Illumina short-read sequencing and short read-only genome assembly was conducted at the DSMZ, as previously described [72]. Hybrid assembly of short and long reads was performed using the Unicycler software (version 0.4.2, bold mode) [71], relying on the programs SPAdes [3] for prior short read assembly, miniasm [27] and Racon [67] for prior long-read assembly and Pilon [68] for polishing of the consensus sequence.

Unless specified otherwise, the following bioinformatic analyses were conducted using the Pantagruel pipeline under the default settings as described previously [23] and on the program webpage http://github.com/flass/pantagruel/. This pipeline is designed for the analysis of bacterial pangenomes, including the inference of a species tree, gene trees, and the detection of horizontal gene transfers (HGT) through species tree/gene tree reconciliations [62]. A more detailed description of genomic datasets and bioinformatic analyses is given in the Supplementary Methods.

### Genomic dataset

We used Prokka [54] as part of *Pantagruel* (task 0) to annotate the new genomes sequences, using a reference database of annotated proteins from the *Rhizobium*/*Agrobacterium* group. We complemented our set of eight new genomes with a dataset of 563 publicly available bacterial genomes obtained from the NCBI RefSeq Assembly database that cover the alphaproteobacterial families *Rhizobiaceae* and (sister group) *Aurantimonadaceae*, for a total of 571 genomes (dataset ‘571Rhizob’).

### Reference species trees

From the 571Rhizob genome dataset, we define the pseudo-core genome (thereafter referred as *pCG*_571_) as the set of genes occurring only in a single copy and present in at least 561 out of the 571 genomes (98%). The *pCG*_571_ gene set includes 155 loci, for which protein alignments were concatenated and used to compute a reference species tree (*S*_ML571_) with RAxML [57] under the model PROTCATLGX; branch supports were estimated by generating 200 rapid bootstraps. From the *S*_ML571_ tree, we identified the well-supported clade grouping 41 genomes including all representative of *Neorhizobium* spp. and *Pseudorhizobium* spp. and our new isolates (dataset ‘41NeoPseudo’). We restricted the *pCG*_571_ concatenated alignment to the 41 genomes of this clade of interest, which we used as input to the Phylobayes program and ran a more accurate (but computationally more expensive) Bayesian phylogenetic inference under the CAT-GTR+G4 model [30] to generate a robust tree for the 41 genomes (*S*_BA41_). We finally used this *S*_BA41_ tree as a fixed input topology for Phylobayes to infer an ultrametric tree (unitless ‘time’ tree) under the CIR clock model [26], further referred to as *T*_BA41_.

### Gene trees, reconciliations and orthologous group classification

Gene trees were computed for each of the 6,714 homologous gene family of the 41-species pangenome with at least four sequences using MrBayes [46] under the GTR+4G+I model. The resulting gene tree samples had the first 25% trees discarded as burn-in and we used the remainder as input for the ALEml program [61,63], to reconcile these gene trees with the reference tree *T*_BA41_ and estimate evolutionary scenarios for each gene family, featuring events of gene duplication, transfer and loss (DTL). Based on the estimated gene family evolutionary scenarios, we could define orthologous gene groups (OGGs) based on a true criterion of orthology, i.e. common descent from an ancestor by means of speciation only [11], rather than a proxy criterion such as bidirectional best hits (BBH) in a similarity search. This has the advantage of explicitly detecting the gain of an OGG in a genome lineage by ways of HGT or gene duplication. We then built a matrix of OGG presence/absence in the ‘41NeoPseudo’ dataset, and computed the clades-specific core genome gene set for each clade of the species tree *S*_BA41_. Hierarchical clustering was performed based on the OGG presence/absence matrix using the pvclust function form the pvclust R package version 2.0-0 [45] with default settings, to obtain bootstrap-derived p-values (BP) and approximately unbiased (AU) branch support estimates. We compared the distribution of functional annotations (Gene Ontology terms) between sets of genes specific to each clade in the *S*_BA41_ tree and corresponding reference gene sets made of the clade’s core-genome or the clade’s pangenome.

### Overall genome relatedness measurement

We used the GGDC tool (version 2.1) for digital DNA-DNA hybridization (dDDH) to compare the genomes of the closest relatives of the NT-25/26 clone, using the formula *d_4_* (BLASTN identities / HSP length) [30]. We also used compareM [37] to estimate the amino-acid average identity (AAI) [20] between genomes of the 41NeoPseudo dataset.

### Biochemical tests

A range of phenotypic assays were performed on a set of ten strains, including the five newly PacBio-sequenced strains as well as the relevant type strains. Salt tolerance: growth of strains was assayed at 28°C in liquid R2A medium (DSMZ medium 830) supplemented with increasing concentrations of NaCl (0–9% range was tested) to determine their minimum inhibitory NaCl concentration (MIC_NaCl_). Congo Red assay: strains were plated on yeast extract – mannitol agar medium (YEM) [44] with 0.1g/L Congo Red dye for seven days. The commercial biochemical identification system for Gram-negative bacteria ‘Api 20 NE’ (BioMérieux) was used for an initial analysis of the biochemical capacities. High-throughput phenotyping was conducted using the GenIII microplates (Biolog, Inc., Hayward, California) for testing for growth with 94 single carbon or nitrogen nutrient sources or with inhibitors (antibiotics, salt, etc.). The GenIII phenotype data were analysed using the ‘opm’ R package [50]. The association of accessory gene occurrence with phenotypic profiles obtained with the Biolog GenIII (continuous values) was tested using the phylogenetic framework implemented in the ‘treeWAS’ R package [7].

### MALDI-TOF typing

Sample preparation for MALDI-TOF mass spectrometry was carried out according to Protocol 3 in [41]. Instrumental conditions for the measurement were used as described by [49]. The dendrogram was created by using the MALDI Biotyper Compass Explorer software (Bruker, Version 4.1.90).

### Fatty acid profiles

Fatty acid methyl esters were obtained as previously described [17] and separated by using a gas chromatograph (model 6890 N; Agilent Technologies). Peaks were automatically computed and assigned using the Microbial Identification software package (MIDI), TSBA40 method, Sherlock version 6.1. The dendrogram was created with Sherlock Version 6.1. Polar lipids and respiratory lipoquinones were extracted from 100 mg freeze-dried cells and separated by two-dimensional silica gel thin layer chromatography by the identification service of the DSMZ as previously described [14].

## Results

### Genome sequencing of eight new *Rhizobiaceae* genomes

We determined the genome sequences of strains *Rhizobium* sp. NT-25, *R. flavum* YW14^T^, *R.* sp. Q54, *R. sp.* TCK and *R.* sp. Khangiran2 using hybrid assembly of Illumina short sequencing reads and PacBio long reads, both at high coverage (Sup. Table S1). Hybrid assembly yielded high-quality complete genomes with all circularized replicons (chromosomes and plasmids) for all strains except Q54. In the genome assembly of strain Q54, only one 463-kb plasmid is circularized, leaving eleven fragments, of which one is chromosomal (size 3.79 Mb) and ten (size range: 1–216 kb) that could not be assigned to a replicon type. In addition, strains AB21, JC140 and P007 were sequenced using Illumina short sequencing reads only; their assembly produced high-quality draft genomes, with 20 to 84 contigs, with N50 statistics ranging 336–778 kb and average coverage 43x–75x (Sup. Table S1). All these genomes carry plasmids, with one to four confirmed circular plasmids in strains NT-25, TCK, YW14 and Khangiran2 (plasmid size range: 15–462 kb) and possibly more for strains Q54, AB21, JC140 and P007.

### Comparison of genomes with digital DNA-DNA hybridization and AAI similarity

To direct the assignment of strain NT-26 and the newly sequenced strains to existing or new species, we proceeded to pairwise comparisons of the new whole genome sequences with the already published reference genomes of strain *R.* sp. NT-26 and type strains of related species *P. pelagicum* R1-200B4^T^ and *R. marinum* MGL06^T^ using dDDH (Table 1). As expected, strains NT-25 and NT-26 are highly related (98% dDDH; 100% AAI) and are thus considered to belong to the same clone (thereafter referred to as the ‘NT-25/26 clone’). They are also closely related to strain TCK (71.5– 71.7% dDDH; 98.4–98.5% AAI), indicating these three strains form a new species. Strains *R.* sp. Q54 groups clearly with *R. endolithicum* JC140^T^ (81% dDDH; 98.2% AAI) and thus is assigned to the species *R. endolithicum*. The dDDH score of 76.30% (97.8% AAI) between the genomes of *P. pelagicum* and *R. marinum* type strains (R1-200B4 and MGL06) indicates that both strains belong to the same species. *R. marinum* [29] having priority over *P. pelagicum* [18,33], this warrants that *R. marinum* should be kept as the valid species name and *P. pelagicum* as its synonym. Strains *R. flavum* YW14^T^, *R. halotolerans* AB21^T^, and *R.* sp. Khangiran2 are closely related, but their dDDH score below the classic 70% species threshold [43], but a AAI score of 97.1% with *R. halotolerans* AB21^T^ (96.5% with *R. flavum* YW14^T^), a value similar to those reported above between species members.

**Table 1.**
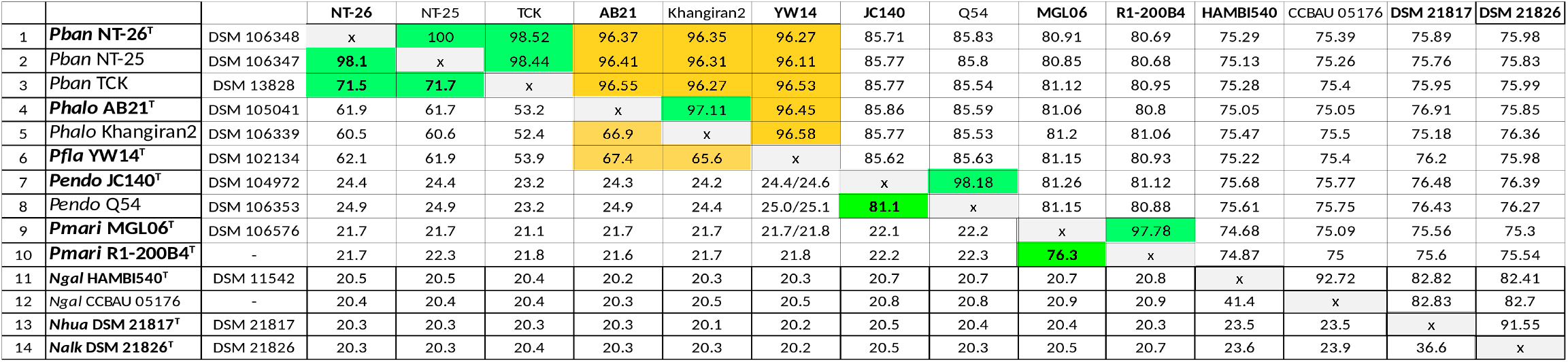
Genome-to-Genome Distance Calculation (GGDC) of similarity between *Pseudorhizobium* strains. Genome similarity estimated using the GGDC tool (version 2.1) with the formula *d_4_* (BLASTN identities / HSP length) to compute dDDH values (lower triangle) and using CompareM to compute AAI values (upper triangle). Similarity values are in percent (%). Green shading indicate values over the 70% dDDH/97% AAI threshold recommended for assignment to the same species; orange shading indicate contentious values close to the threshold.

While dDDH value saturate around 20% when comparing distant species (Table 1), AAI values decrease more gradually. AAI values between type strains from different genera do not exceed 76%, whereas values between types of *Neorhizobium* species are all above 82%, suggesting a discontinuity of AAI values can be used to determine genus membership. Interestingly, the strain cluster including NT-26 and the type strains of *R. flavum, R. halotolerans*, *R. endolithicum*, *R. marinum* and the type strain of the *Pseudorhizobium* genus, R1-200B4^T^ (= LMG 28314^T^ = CECT 8629^T^), all show AAI values above 80%, suggesting membership of a same genus (Table S2). The genus *Rhizobium* is well known to be paraphyletic and several new genera have been recently defined to solve that issue [31,32]; our observation further the reclassification of *R. halotolerans, R. flavum*, *R. endolithicum* and *R. marinum* into the *Pseudorhizobium* genus, thus becoming *P. halotolerans* (*Phalo*)*, P. flavum* (*Pfla*), *P. endolithicum* (*Pendo*) and *P. marinum* (*Pmari*).

### Phylogeny of *Neorhizobium* and *Pseudorhizobium*

We produced a large phylogeny based on the concatenated 155 core proteins of 571 *Rhizobiaceae* and *Aurantimonadaceae* complete genomes and using a fast ML method of inference (*S*_ML571_) (Sup. Fig. S1). In addition, we generated a phylogeny focused on the group of interest encompassing the *Neorhizobium* and *Pseudorhizobium* genera (‘41NeoPseudo’ dataset) using a Bayesian inference and more realistic molecular evolution model (*S*_BA41_) (Figure 1), which confirmed the groupings described above based on dDDH and AAI. Almost all branches in *S*_BA41_ are well supported with Bayesian posterior probability (PP) support > 0.97, except some internal branches in the *Neorhizobium* clade and the branch grouping strains *R.* sp. Khangiran2, *Phalo* AB21^T^ and *Pfla* YW14^T^. Both the *S*_ML571_ and *S*_BA41_ trees place strain Khangiran2 closer to AB21^T^ than to YW14^T^, in agreement with pairwise AAI values. This indicates strain Khangiran2 should be classified as a member of *P. halotolerans.* The relatively low PP support of 0.79 on the stem branch of the *Pfla+Phalo* group suggests that gene flow may have occured between this group and theirclose relatives.

**Figure 1:**
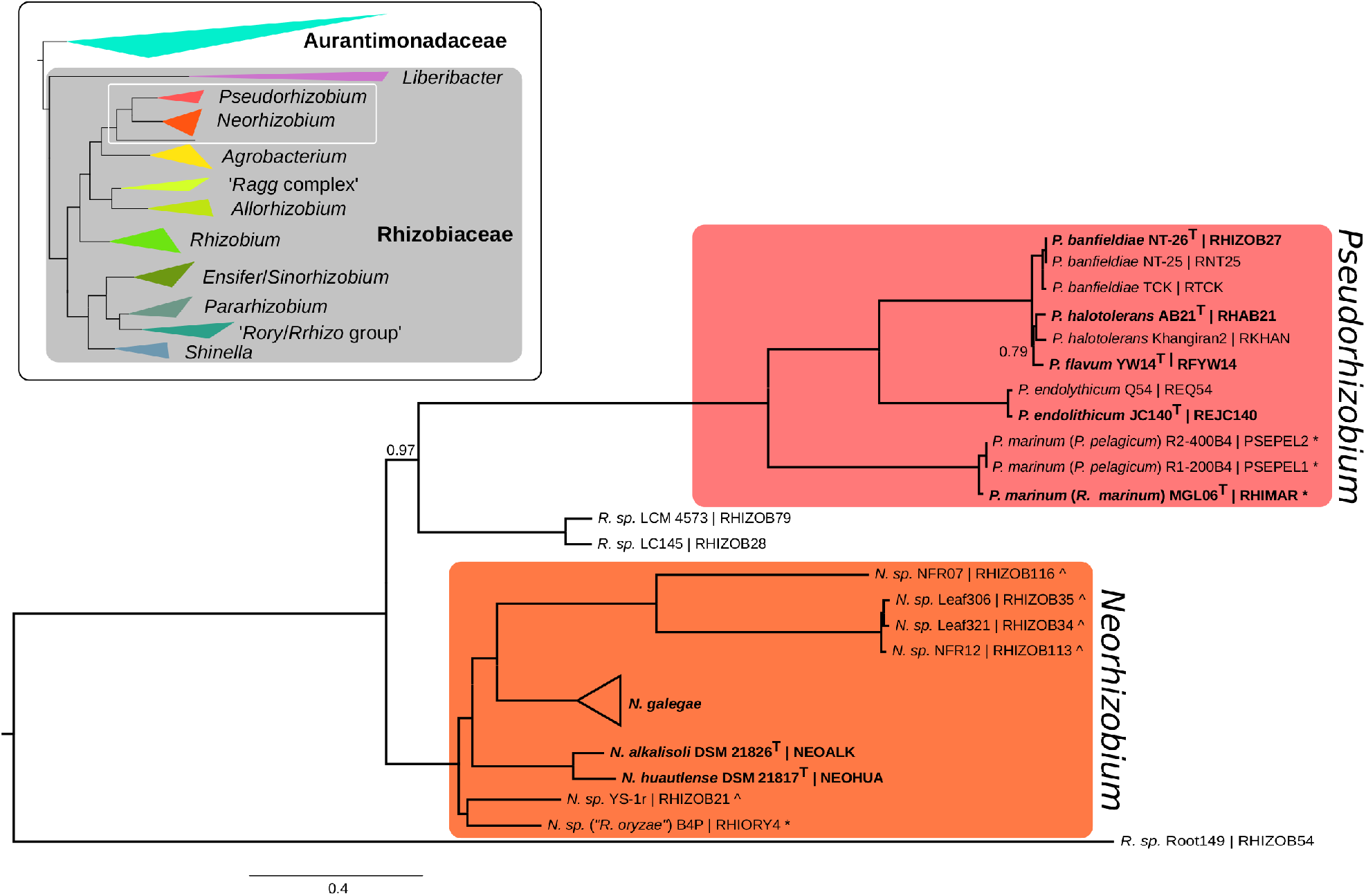
Bayesian phylogenetic tree of 41 organisms from the *Neorhizobium* and *Pseudorhizobium* genera and close relatives (*S*_BA41_). Tree obtained with Phylobayes under the GTR-CAT protein evolution model, based on a concatenated alignment of 155 pseudo-core protein loci. All posterior probability branch supports are 1.00 unless indicated. The organism name is followed (after the pipe symbol |) by the identifier in the Pantagruel pangenome database of this study. Strains whose species or genus affiliation are corrected in this study are marked with an asterisk * or caret ^, respectively. Species type strains are in bold. The clade *N. galegae* was collapsed; it includes the type strain *N. galegae bv. orientalis* HAMBI 540^T^ (organism id: NEOGAL2). The full tree is presented in Sup. Fig. S2. A schematic view of the phylogenetic context of this group is indicated in inset (based on tree *S*_ML571_, which presented in full in Sup. Fig. S1). *Raag*: *R. aggregatum*; *Rory*: *R. oryzae*; *Rrhizo*: *R. rhizosphereae*. The alignment and tree files are available on Figshare (doi: 10.6084/m9.figshare.8316827).

The clade containing strains NT-25, NT-26 and TCK groups with the *Phalo+Pfla* clade, and further with *Pendo* and then *Pmari*, as a well-separated clade from *Neorhizobium*, i.e. the *Pseusdorhizobium* genus. Accordingly, we propose that strains NT-25, NT-26 and TCK should form a new species in this genus and we propose to name it “*Pseudorhizobium banfieldiae*” (*Pban*).

The positions of strains “*Rhizobium oryzae*” B4P and “*R. vignae*” CCBAU 05176 in the *S*_ML571_ and *S*_BA41_ phylogenetic trees (Sup. Fig. S1, S2) and their pairwise AAI values (Table S2) makes it clear that they were incorrectly named and should be designated as *Neorhizobium.* sp. B4P and *N. galegae* CCBAU 05176. Similarly, other strains present in the clade corresponding to the genus *Neorhizobium* and currently identified as *Rhizobium* sp. need to be renamed as *Neorhizobium* sp.: strains YS-1r, NFR07, NFR12, Leaf306 and Leaf321.

We compared the phylogenies based on core genome alignment to those obtained with alternative sources of information on genomic variation (detailed in Suppl. Text). Based on the distribution of pangenome genes, i.e. the presence/absence of orthologous accessory genes in the ‘41NeoPseudo’ genomes (*S*_CL41_), a hierarchical clustering shows a very similar picture to *S*_BA41_, with good support for most branches leading to major clades and species (Figure 2B; Sup. Fig. S3). Low support for the branch separating *Pfla* YW14^T^ from *Phalo* strains Khangiran2 and AB21^T^ suggests frequent HGT between these species. Additionally, strains that branch deep in the *S*_BA41_ tree all cluster together as a sister group of the *Pseudorhizobium* clade. This limited resolution of gene presence/absence data beyond the species level may be explained by inter-specific HGT, possibly driven by convergent adaptation.

**Figure 2:**
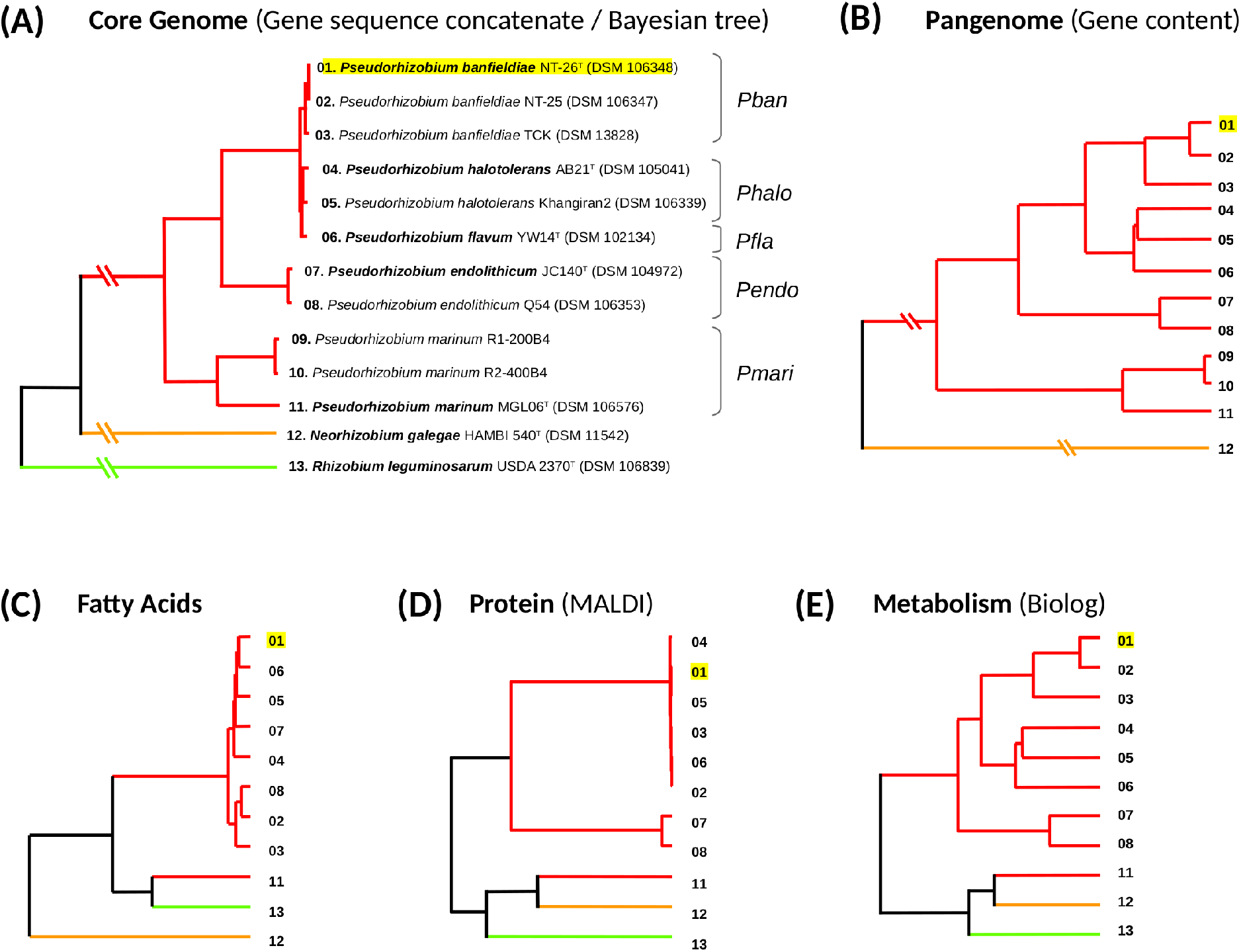
Phylogenetic clustering of Pseudorhizobium strains based on genomic and phenotypic characters. A numeric code corresponding to strains as indicated in panel (A) is used in the other panels. **A.** Bayesian tree *S*_BA41_ based on core genome gene concatenate, adapted from Figure 1; *R. leguminosarum* is indicated as an outgroup, in accordance with the maximum-likelihood tree *S*_ML571_ based on the same core genome gene set and wider taxon sample. **B**. Hierarchical clustering dendrogram based on the accessory gene content of strains defined at the orthologous group level. **C-E**. Hierarchical clustering dendrogram based on phenotypic data relating to fatty acid content, protein content and metabolic abilities. Underlying data are available on Figshare: doi: 10.6084/m9.figshare.8316827; doi: 10.6084/m9.figshare.8316383; doi: 10.6084/m9.figshare.8316746.

Phylogenetic trees were also built from a restricted set of classic marker genes (*atpD*, *recA*, *rpoB, glnII* and *gyrB*), either separately or in concatenation, i.e. in a multi-locus sequence analysis (MLSA) (Supp. Methods; Supp. Dataset S1). The monophyly of all species within *Pseudorhizobium* is recovered with the MLSA, but not with any single marker gene. However, the deeper grouping *Phalo*+*Pfla* is not recovered by the MLSA tree, with the inclusion of *Pban* in that clade being moderately supported (see Suppl. Text). These results indicate that marker gene-based analyses are mostly consistent with the information obtained from whole genomes, with MLSA providing a satisfying framework for species typing of *Pseudorhizobium* strains. However, the study of deeper evolutionary relationship and the classification of distant strains should be based on genome-wide data, in accordance to recent guidelines [22].

### Phenotype-based classification

We tested the ability of high-throughput phenotyping methods to generate taxonomic information. We found that clustering of strains based on data generated from lipid, protein or metabolic screens yielded a classification broadly similar to the one obtained from core-genome alignments. Fatty acid profiling showed the poorest resolution as it could not discriminate between species (Figure 2C). However, it distinctly clustered *Pban*+*Phalo*+*Pfla*+*Pendo* to the exclusion of its outgroups *Neorhizobium* and *P. marinum*, indicating a synapomorphic change of lipid composition at the common origin of these four species. A proteome screen (using MALTI-TOF mass spectrometry) further differentiated *Pendo* from the group *Phalo*+*Pfla*+*Pban* (Figure 2D). Metabolism profiles (based on growth curves in 95 different conditions) proved the most accurate sequence-independent predictor of the phylogenomic tree as it also distinguished the group *Phalo*+*Pfla* from *Pban* and thus almost completely mirrored the branching pattern in the genus *Pseudorhizobium* (Figure 2E). All phenotype screens however led to cluster *Pmari* MGL06^T^ with outgroups *N. galegae* and *R. leguminosarum* (Figure 2C-E), showing the limited ability of chemotaxonomic and phenotypic analyses to resolve taxonomy at deeper evolutionary scales, likely due to convergence of adaptive traits.

### Clade-specific gene sets reveal specific functions and ecologies

We inferred gene family evolution scenarios accounting for HGT history by reconciling gene tree topologies with that of the species tree *S*_BA41_. Based on these scenarios, we delineated groups of orthologous genes that reflect the history of gene acquisition in genome lineages – every gain of a new gene copy in a genome lineage creating a new OGG. We looked for groups of OGGs with contrasting occurrence patterns between a focal clade and its relatives, to identify specific events of gene gain or loss that led to the genomic differentiation of the clade. Data for all clade comparisons in our ‘41NeoPseudo’ dataset are presented in Sup. Table S3 and are summarized below for the clades on the lineage of strain NT-26; more detailed information and description of gene sets specific to other groups are listed in the Supplementary Text. Major gene sets that have contrasting pattern of occurrence in *Pseudorhizobium* are listed Table 2 and those specifically contributing to the differentiation of the NT-25/26 clone lineage are depicted in Figure 3.

**Table 2.**
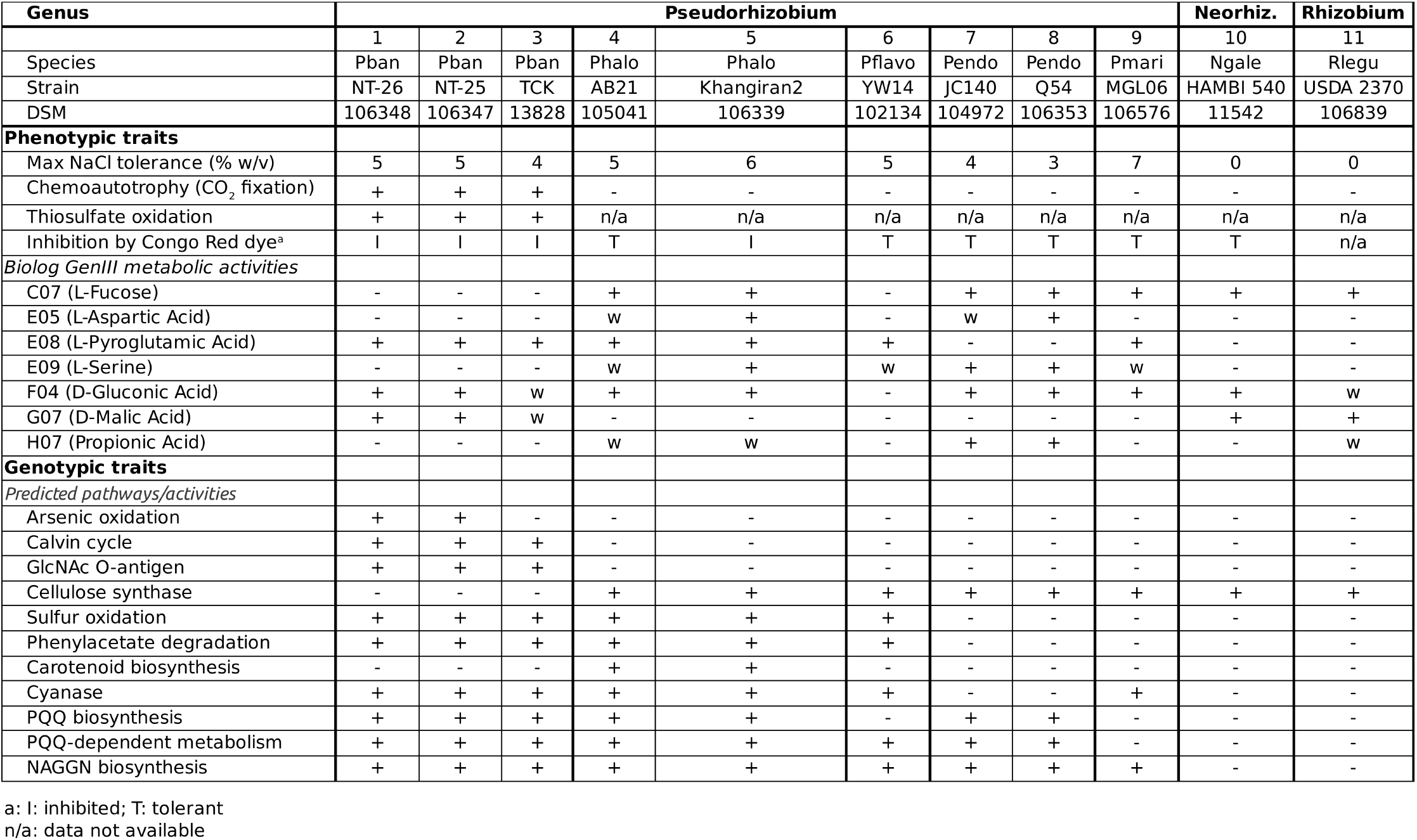
Main Phenotypic and Genotypic features distinguishing Pseudorhizobium species.

**Figure 3:**
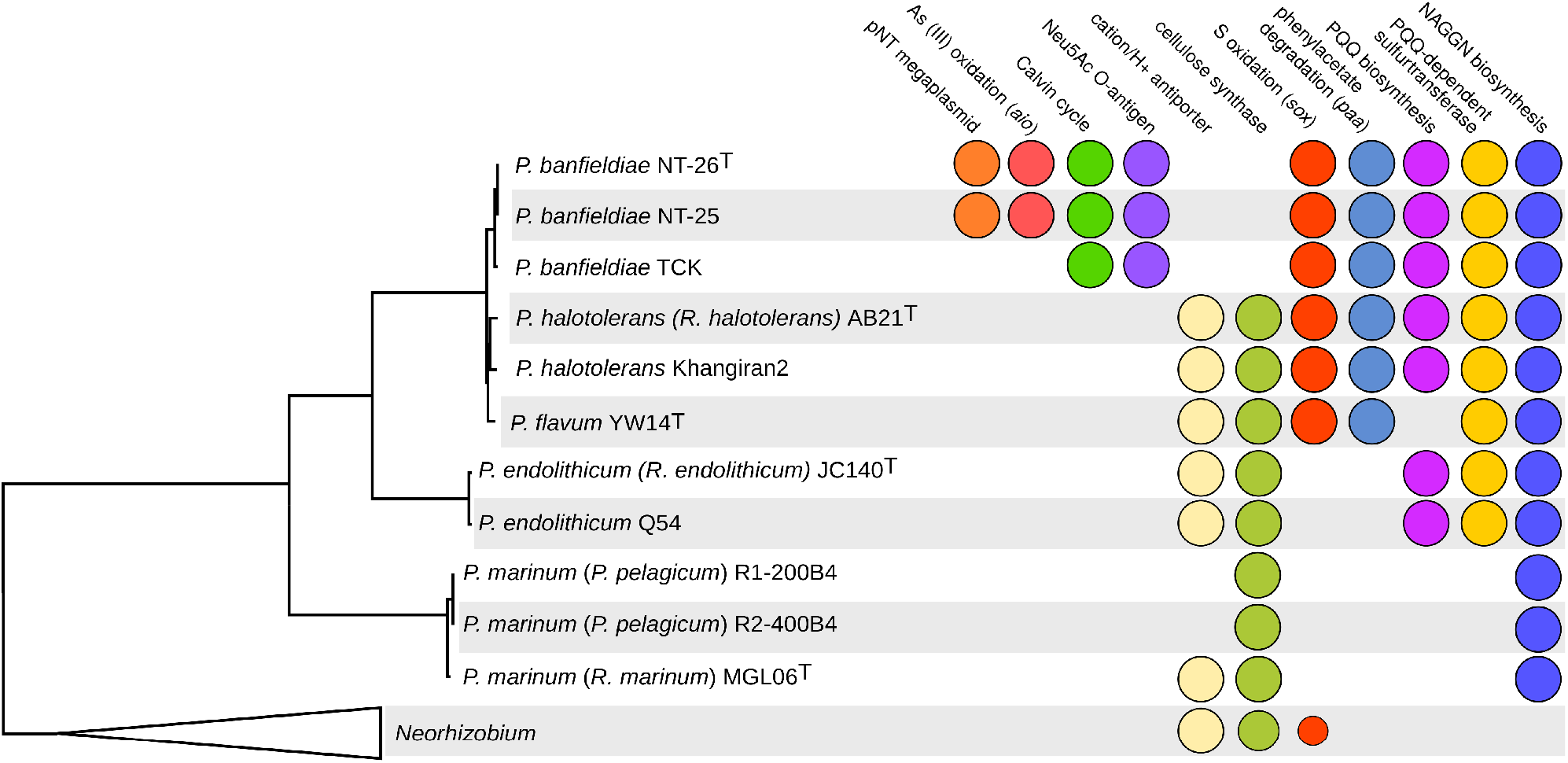
phylogenetic distribution of selected pathways in *Pseudorhizobium*. A circle indicates the presence of the genes or pathway in the genome. For the collapsed *Neorhizobium* clade, the frequency of presence of the genes is indicated by the area of the circle.

#### NT-25/NT-26 clone

Most of the gene content specific to this group is composed of mobile or selfish elements. As expected, this includes the 322-kb plasmid in NT-26 and homologous 119-kb plasmid in NT-25 that carry the arsenite oxidation *aio* locus and extra copies of the arsenic resistance *ars* operon and the phosphate-specific transporter *pst*/*phn* locu*s*. In addition, the chromosome is laden with specific mobile elements. A prophage is located between two tRNA genes (positions 1,347–1,419 kb; length 71 kb) characterized by an entire set of phage structural genes and an integrase gene at the end of the locus, with no other identified function in its gene cargo. A putative integron (positions 352–394kb; length 41 kb) is characterized by an integrase gene at the end of the locus, next to a tRNA gene; it also carries several genes involved in transport and metabolism of a putative branched-chain carbohydrate substrate.

#### ‘Pseudorhizobium banfieldiae’ (Pban)

Of all the genes exclusively present in *Pban* compared to other *Pseudorhizobium*, the most striking feature is a locus encoding the Calvin cycle path way (including the RuBisCO enzyme) and respiratory chain cytochromes, the main determinants of chemoautotrophy in this species. This locus belongs to a larger *Pban*-specific region composed of two closely located 27-kb and 80-kb fragments, which suggests it results from the recent insertion and domestication of a mobile element (likely interrupted by an even more recent insertion/rearrangement). Among the 37 *Pban*-specific genes in this extended region, several code for enzymes of the classes oxidoreductase, monooxygenase, decarboxylase and glutathione-S transferase, which all use reduced electron acceptors and/or protons, and with their putative substrates including aromatic cyclic and halogenated organic compounds.

This suggests a functional link between chemoautotrophy and detoxification pathways. Reconstruced HGT scenarios indicate that the donor of these genes was a deep-branching lineage of *Neorhizobium* (Sup. Fig. S4; Sup. Table S4), but also that it was preceded by series of HGT events, dated as early as the diversification of the *Neorhizobium/Pseudorhizobium* group. This suggests genes coding for chemoautotrophy have been circulating for a long time in this wider taxon, and were later fixed in the *Pban* lineage. Other *Pban-*specific genes include a locus putatively encoding the biosynthesis of a lipopolysaccharide O-antigen with an N-acetylneuraminic acid (Neu5Ac) function, and a 26-kb region encoding putative enzymes and transporters related to pathways for the utilization of taurine and for the degradation of (possibly halogenated) aromatic compounds (Sup. Table S5).

Conversely, several genomic regions have been lost in the *Pban* lineage. An operon that encodes a multimeric Na^+^/H^+^ cation antiporter was present in the ancestor of of *Pban*, *Phalo, Pfla* and *Pendo*, then specifically lost in *Pban*; an homolog is present in *Pmari* strain MGL06^T^, with the gene evolution scenario indicating this gene is a HGT recipient from *Phalo* to MGL06. An operon encoding a cellulose synthase is present in all other *Pseudorhizobium* species, indicating the likely presence of a cellulose-like polymer in their exopolysaccharide, but not in *Pban* where it was lost. Finally, *Pban* genomes specifically lack genes coding for a respiratory complex including several cytochrome *c* oxidases, in linkage with a gene encoding the EutK carboxysome-like microcompartment protein, whose known homologues are involved in the degradation of ethanolamine (see Supplementary text). This respiratory complex-encoding locus often includes genes coding for redox enzymes that may be the terminal electron acceptor; interestingly, these genes vary with the species (Sup. Fig. S5): *Pfla* YW14 carries a copper-containing nitrite reductase, while *Phalo* strains AB21 and Khangiran2 carry a (non-homologous) TAT-dependent nitrous-oxide reductase; the locus in *Pendo* strains harbours no gene encoding such terminal electron acceptor, but other genes encoding metabolic enzymes that differ among strains. This suggests that this respiratory chain and associated putative micro-compartment are used as an evolutionary flexible platform for the reductive activities of these organisms.

#### Pseudorhizobium sub-clade Pban+Phalo+Pfla

The *Pban*+*Phalo+Pfla* clade presents two large specific gene sets: the 20-kb super-operon *paa* coding for the uptake and degradation of phenylacetate, and the 13-kb locus including the *soxXYZABCD* operon that encodes the sulphur oxidation pathway, allowing the lithotrophic oxidation of thiosulphate.

#### Pseudorhizobium sub-clade Pban+Phalo+Pfla+Pendo

Genes specific to the *Pban*+*Phalo+Pfla+Pendo* clade are enriched in the cellular process of NAD cofactor biosynthesis (GO:0009435), tryptophan catabolism (GO:0019441) and phosphatidic acid biosynthesis (GO:0006654). They also carry an operon encoding a thiosulphate sulphurtransferase with a pyrroloquinoline quinone (PQQ)-binding motif, a SoxYZ-like thiosulphate carrier and a SoxH-like metallo-protease and a membrane-bound PQQ-dependent dehydrogenase with glucose, quinate or shikimate as predicted substrates. In addition, a 17-kb locus including the *pqqBCDE* operon involved in the biosynthesis of cofactor PQQ and PQQ-dependent methanol metabolism enzymes was also specifically gained in this clade, but later lost by *Pfla* strain YW14.

#### Pseudorhizobium (Pban+Phalo+Pfla+Pendo+Pmari)

In comparison with the closely related genus *Neorhizobium, Pseudorhizobium*-specific genes are over-represented in genes involved in cellular processes related to energy metabolism: ‘aerobic respiration’ (GO:0009060) and ‘electron transport coupled proton transport’ (GO:0015990), and to anabolic processes, including the biosynthesis of cofactor NAD (GO:0009435), lipid precursor acetyl-CoA from acetate (GO:0019427) and amino-acid asparagine (GO:0006529). In addition, a *Pseudorhizobium*-specific operon encodes the biosynthesis of osmoprotectant N-acetylglutaminylglutamine (NAGGN).

#### Other Pseudorhizobium species

Traits specific to other clades, including the species *Pendo*, *Pmari* and *Phalo*, are discussed in the Supplementary Text. Among the many species-specific traits found, we can highlight the following predictions: *Phalo* features a specific pathway involved in the biosynthesis of carotenoids; *Pendo* has specific accessory components of its flagellum, and misses many genes that are otherwise conserved in the genus, including a cyanase gene; *Pmari*, as the most diverged species in the genus, has several hundred species-specific genes, including a 27-kb locus coding for a potassium-transporting ATPase, extrusion transporters and degradation enzymes with putative phenolic compound substrates, and a poly(3-hydroxybutyrate) (PHB) depolymerase.

### Validation of bioinformatic predictions of phenotypes

We aimed to experimentally validate the predictions of clade-specific phenotypes that would allow us to distinguish taxa and also to confirm the bioinformatically predicted functions of the identified genes. We thus implement a new version of the polyphasic approach to taxonomy [43], where genome-based discovery of phenotypes complements genome relatedness-based delineation of taxa, and ultimately would help us link the conservation of a genotype in a taxon with relevant aspects of its ecology [24]. We focus on *P. banfieldiae*, for which our dataset provides the best phylogenetic contrast, with three genomes sampled within the taxon and eight genomes sampled in close relatives, ensuring the robust identification of species-specific genes (Figure 3). We describe below the clade-specific phenotypic traits that were experimentally validated. For other predicted traits, i.e. the specific utilization of taurine and phenylacetate for *Pban* and *Pban*+*Phalo*+*Pfla* clades, respectively, the experimental test did not match the expectations (see Supplementary Text).

Pban*-specific chemolithoautotrophy.* This is a known trait of all *Pban* strains, which were indeed isolated for that particular property [9,40]. All strains can grow with thiosulphate as a sole electron source and by fixing carbon dioxide as a C source; the use of arsenite as an electron donor is unique to the NT-25/26 clone, due to the presence of the *aio* operon on the clone’s specific plasmid.

*Genus-wide salt tolerance.* Tolerance of salt is a known trait of all previously reported strains of *Phalo* (up to 4% NaCl)*, Pfla* (up to 4% NaCl)*, Pendo* (up to 5% NaCl), and *Pmari* (up to 7% NaCl for former *P. pelagicum* strains and up to 9% NaCl for strain MGL06^T^), having indeed inspired the choice of the species epithet of *R. halotolerans* [10,13,18,29,36]. This phenotype could be conferred, at least in part, by the expression of a Na^+^/H^+^ antiporter, a function that was identified as a marine niche-associated trait in *Rhodobacteraceae* [42]. The Na^+^/H^+^ antiporter genes are missing in *Pban*, leading us to predict a lower salt tolerance in this species (Figure 3). The ability to grow in NaCl concentrations ranging from 0 to 9% was tested for 11 strains of *Pseudorhizobium* and related organisms (Sup. Table S6). The results showed no significant difference between *Pban* and other species in the genus with all tolerating up to 3–6% NaCl under our test conditions, apart from *Pmari* strain MGL06, which still grew in the presence of 7% NaCl, rejecting the hypothesis of the presence of a Na^+^/H^+^ antiporter as cause of this phenotype. The common baseline of salt tolerance in *Pseudorhizobium* suggests that core genes encode salt tolerance factors or, less parsimoniously, that all lineages have convergently evolved such traits. The levels of salt tolerance we measured are lower than previously reported for *Pendo* and *Pmari*, and higher for *Phalo*, suggesting that other factors that determine salt tolerance in other conditions were not expressed in our experiment.

Pban*-specific lack of production of a cellulose polymer. Pban* and *Phalo* strains were plated on yeast extract–mannitol agar medium (YEM) supplemented with 0.1g/l Congo Red dye, a characteristic marker of beta-glucan polymers, to test for the presence of cellulose or a related polymer such as curdlan in their capsular polysaccharide [19]. Contrary to expectations, *P. banfieldiae* strains were coloured by the dye, with an orange-red hue, which was observed for other tested *Pseudorhizobium* strains (Sup. Fig S6; Sup. Table S7). However, growth of *Pban* strains was inhibited in the presence of the Congo Red dye, resulting in small, dry colonies on YEM plates (Sup. Fig S6; Sup. Table S7) or no growth on MSM + 0.08% YE (data not shown). This growth phenotype was unique to *Pban* among *Pseudorhizobium*, except for *Phalo* strain Khangiran2 (which was coloured salmon-orange). The inhibitory effect of the dye has been previously observed for other bacteria deficient in the production of beta-glucan polymers [46], and the inhibition observed on *Pban* strains could thus reflect the absence of protection that the cellulose-like polysaccharide provides to other *Pseudorhizobium* isolates. The case of strain Khangiran2 remains unclear as it seems to strongly bind the dye (thus suggesting the polymer is expressed), but it nonetheless suffers from exposure to the dye.

### Search for genotype-phenotype associations

We took advantage of our well-defined phylogenomic framework and of the compendium of phenotypes that we had tested (Sup. Table S8) to search for potential associations between the distribution of accessory genes and the distribution of phenotypes, with the expectation of revealing new gene functions. Using a genome-wide association (GWAS) testing framework, we looked for the basis of significant phenotypes that are not necessarily distributed following the taxonomical structure of species. Specifically, we explored the association between metabolic traits and the distribution of OGGs in the accessory genome, using the species tree to account for potential spurious associations linked to oversampling of closely related strains. The GWAS reported numerous significant associations, listed in Sup. Table S9. Manual exploration of results singled out only one association with a clear association pattern that we believe to be of biological relevance: the utilization of beta-methyl-D-glucoside, observed most strongly in strains *Pban* NT-26, *Phalo* AB21 and *N. galegae* HAMBI 540, was associated with the presence of several chromosomal clusters of genes. These include two operons, one encoding an allophanate hydrolase, and another encoding a transporter and a dienelactone hydrolase-related enzyme.

## Discussion

The traditional polyphasic approach in bacterial taxonomy combines several criteria, in particular marker gene phylogeny and biochemical phenotypes, to determine the boundaries of taxa [43]. The validity of this approach has recently been questioned, due to the growing evidence that most phenotypes are encoded by genes that may be accessory within a species or be shared promiscuously among species, making them poor diagnostic characters [21]. Instead, the growing practice in the field is to use whole genome sequences to estimate similarity between bacterial isolates [5] and to compute phylogenetic trees based on genome-wide data [34]. A tree provides the hierarchical relationships between organisms, while the level of overall genome relatedness provides an objective criterion for the delineation of species. This criterion requires a threshold, and 70% has been proposed for dDDH, which is equivalent to the classical score of DNA-DNA hybridization (DDH) widely considered as adequate for species delineation, [4], as obtained using the GGDC tool [30]; we also established that in this dataset, 97% amino-acid average identity (AAI) provides a practical species threshold. However, similarity thresholds are arbitrary and the relevance of species boundaries proposed based on this sole criterion may be questioned.

In this study, pairwise genome similarities among strains Khangiran2, *Phalo* AB21^T^ and *Pfla* YW14^T^ are all below, but close to the thresholds of 70% dDDH and 97% AAI. According to recent taxonomical guidelines [22], criteria other than similarity-based ones should be considered to decide on species boundaries. We therefore complemented our taxonomic investigations with an assessment of the genomic, chemotaxonomical and ecological differentiation between strains. These three strains form a clade (‘*Phalo+Pfla*’) that is well supported in the *S*_ML571_ ML tree. Genes specific to this clade are enriched in functions involved in the biogenesis and modification of membrane lipids. This important cellular pathway could be the means of adaptation to a shared ecological niche, and this group could thus constitute an ecological species [24]. However, core genes specific to *Phalo* (i.e. strains AB21 and Khangiran2) are also significantly enriched with coherent cellular functions, including the biosynthesis of carotenoid pigments and related isoprenoid metabolism – a key metabolic pathway suggesting that the core genome of *Phalo* may also be involved in the adaptation to its own specific niche. Therefore, the ecological arguments rather support *Phalo* and *Pfla* to be two distinct ecological species. In addition, the relatively lower support in *S_BA4_*_1_ Bayesian tree for the *Phalo+Pfla* clade is a strong argument against its election to species status. For this reason, we recommend that *P*. *halotolerans* and *P. flavum* shall remain two distinct species until further evidence to the contrary.

While a recent large-scale analysis of *Alphaproteobacteria* type strain genomes suggested that the genus *Pseudorhizobium* should be merged into *Rhizobium* [16], we bring extensive evidence that *Pseudorhizobium* is well differentiated genomically and phenotypically and forms a *bona fide* bacterial genus. Through the phylogeny-aware comparison of genomes, we explored the functional specificities of lineages within this genus (Figure 3). From the functional annotation of genomes, we identified the genomic basis of known phenotypic traits, such as chemolithoautotrophy or sulfur oxidation, and predicted others such as a cellulose component of the bacterial coat or a lipopolysaccharide O-antigen. We mapped the distribution of these traits within a phylogenetic framework, identifying those traits which presence or absence was exclusive to a group. The only new prediction of contrasting phenotypes that could be verified in the lab is related to the absence of a cellulose-like polymer in *Pban*. Phenotypic features of wider evolutionary groups were documented, including the general tolerance of members of the *Pseudorhizobium* genus to NaCl (Sup. Table S6; Supplementary Text). This shared feature suggests that the ancestor of the group might have been itself salt tolerant, and thus possibly a marine organism – a hypothesis consistent with the basal position in the genus tree of seaborne *P. marinum*.

We showed that some cellular processes and pathways were over-represented in the specific core genome of clades of the *Pseudorhizobium* genus. This pattern results from the serial acquisition of genes with related functions in a ancestral lineage and their subsequent conservation in all descendants – a process likely driven by positive selection [25]. Indeed, under an ecotype diversification model, the acquisition of genes enabling the adaptation to a different ecological niche can trigger the emergence of a new ecotype lineage [6]. Ecological isolation of this ecotype may in turn drive the differentiation of its core genome, with additional adaptive mutations (including new gene gains and losses) producing a knock-on effect leading to ecological specialization [24]. Analysing the specific gene repertoire of each clade of the *Pseudorhizobium* genus indeed shed light on putative ways of adaptations of these groups to their respective ecological niches.

The emergence of the highly specialized NT-25/NT-26 clone could be explained by a hypothetical scenario involving a sequence of ecotype diversification events: a first key event was the acquisition of multiple new cytochromes and interacting redox enzymes in the *Pseudorhizobium* genus ancestor, enhancing its capacity to exploit the redox gradients between available environmental compounds. This was followed by the acquisition by the ancestor of *Pban*+*Pfla*+*Phalo*+*Pendo* of a first set of sulphur oxidation enzymes, with electrons from the periplasmic oxidation of thiosulphate being transferred to the carrier molecules SoxYZ and PQQ, likely to fuel oxidative enzymes such as a jointly acquired toxic carbohydrate-degrading metallo-hydrolase. Then, the *sox* gene cluster was gained by the ancestor of *Pban*+*Pfla*+*Phalo*, allowing it to use thiosulphate as a source of electrons to fuel respiration and therefore to convert them into proton motive force and to recycle the cellular pool of redox cofactors. The joint acquisition of the phenylacetate degradation pathway – many reactions of which require reduced or oxidized cofactors [48] – allowed this organism to use this aromatic compound and its breakdown products as carbon and electron source. This set of new abilities would have allowed this lineage to colonize new habitats either depleted in organic nutrients or contaminated with toxic organic compounds. This was followed by the acquisition of RuBisCO and other Calvin cycle genes and additional respiratory cytochromes by the *Pban* ancestor. The encoded metabolic pathways provide electrons to the cell and allow the fixation of carbon dioxide, thus allowing that ancestor to live chemolithoautotrophically using sulphur oxidation – again this has likely let this lineage colonize environments yet uncharted by most rhizobia, such as rock surfaces. Finally, the acquisition of a plasmid carrying the arsenite oxidation genes and other factors of resistance to arsenic and heavy metals, allowed the NT-25/NT-26 clone ancestor to successfully colonize the extremely toxic and organic nutrient-poor environment of a gold mine.

Aside from this scenario of extreme specialization towards chemolithoautotrophy and resistance against toxic heavy metals, all species in the genus *Pseudorhizobium* have achieved significant ecological differentiation from the *bona fide* rhizobial lifestyle, which is characteristic for members of the most closely related genus *Neorhizobium* typically isolated from soil and the plant rhizosphere [15,28,32,35,51]. *P. endolithicum* has only been found inside the mineral matrix of sand grains and its high salt and temperature tolerance indicate it is likely adapted to this peculiar lifestyle, even though its capacity to nodulate soybean indicates its ecological niche encompasses various lifestyles [36]. Among the many gene gains and losses that occurred over the long branch leading to *Pendo*, one notable change involved the structural and biosynthetic genes of the flagellum, possibly leading to a deviant morphology of this bacterial motor in this species. This might be linked to its ability to colonize the interior of sand rock particles necessitating a particular type of motility.

The *P. marinum* core genome is largely differentiated from the rest of the genus, owing to its early divergence. Among its species-specific components, some genes are involved in functions that are key for survival in a typically marine lifestyle. These include transport of K^+^ and Cl^−^ ions, urea and various sugars and organic acids and amino-acids and the degradation of toxic phenolic compounds, in combination with many signal transduction systems, which must allow the rapid scavenging/extrusion of rare/excess ions or toxins in response to changing availability of mineral and organic nutrients and the rise in toxicity of the environment. In addition, the (de)polymerisation of storage compound PHB may allow the cell to survive long-term starvation during nutrient-depleted phases. Finally, the ability to synthesize of the osmoprotectant NAGGN – a trait common to all sequenced members of the genus *Pseudorhizobium* – makes this species particularly adapted to life in marine habitats and other environments where salinity can vary strongly.

## Conclusion

In summary, in this work we have used a comparative genomics approach within a phylogenomic framework to identify the unique characters of five species of the genus *Pseudorhizobium*, shedding light on the genome evolution that led them to adapt to their respective ecological niche. Our analysis highlights how species – and higher groups – within this clade of the *Rhizobiaceae* family evolved towards strikingly different ecological strategies, through the acquisition of traits such as tropism and resistance to environmental toxins, thus allowing each species to colonize its own peculiar niche.

### Emended description of the genus *Pseudorhizobium* Kimes *et al.* 2015

*Pseudorhizobium* (Gr. adj. *pseudes* false; N.L. masc. n. *Pseudorhizobium*, false *Rhizobium*).

Aerobic, Gram negative non-spore forming rods forming white colonies on YMA. Optimal growth at 28–30°C and pH7-8. Catalase test is positive. Production of β-galactosidase is positive. The production of indol is negative. The production of arginine dehydrolase and gelatinase were negative. Growth was observed in presence of 0–4% NaCl, and up to 7% for certain isolates. The main fatty acids are C_18:1_ω7c/C_18:1_ω6c. The G + C content of genomic DNA is 61.8–62.8 mol%.

The type species is *P. marinum* (synonym: *P. pelagicum*) and the type strain is *P. marinum* R1-200B4^T^.

Delineation of the genus was determined based on whole-proteome similarity analysis (AAI) and the phylogenetic analysis of the concatenated alignments of 155 conserved genes. Strains within the genus have all above 80% AAI similarity between each other, and below 76% AAI similarity with *N. galegae* HAMBI 540^T^, the type strain of sister genus *Neorhizobium*.

### Description of *Pseudorhizobium halotolerans* sp. nov

The description of the species is the same as the descriptions given by Diange and Lee [10], except that it is tolerant to NaCl up to 5 % (w/v), instead of 4 %.

Basonym: *Rhizobium halotolerans* Diange and Lee, 2013 (effectively, but not validly published).

The type strain, AB21^T^ (= DSM 105041^T^ = KEMC 224-056^T^ = JCM 17536^T^), was isolated from chloroethylene-contaminated soil from Suwon, South Korea.

### Description of *Pseudorhizobium flavum* comb. nov

The description of the species is the same as the descriptions given by Gu *et al. [13]*. Notably, it has a tolerance of 0–4 % NaCl (w/v).

Basonym: *Rhizobium flavum* Gu *et al.* 2014.

The type strain, YW14^T^ (= DSM 102134^T^ = CCTCC AB2013042^T^ = KACC 17222^T^) was isolated from organophosphorus (OP) insecticide-contaminated soil.

### Description of *Pseudorhizobium endolithicum* comb. nov

The description of the species is the same as the descriptions given by Parag *et al.* [36]. Basonym: *Rhizobium endolithicum* Parag *et al.* 2013.

The type strain is JC140^T^ (= DSM 104972^T^ = KCTC32077^T^ = CCUG64352^T^ = MTCC11723^T^ = HAMBI 2447^T^), isolated from sand rock matrix.

### Description of *Pseudorhizobium marinum* comb. nov

The genus *Pseudorhizobium* was described along with the species name *P. pelagicum* Kimes *et al.* 2015, which is a heterotypic synonym of *R. marinum* Liu *et al.* 2015. Because the genus name *Pseudorhizobium* remains valid (with type strain R1-200B4^T^ = LMG 28314^T^ = CECT 8629^T^) [33], the species epithet *marinum* is now to be preceded by the *Pseudorhizobium* genus prefix.

The description of the species is the same as the descriptions given by Liu *et al.* [29].

Basonym: *Rhizobium marinum* Liu *et al.* 2015

The type strain is MGL06^T^ (= DSM 106576^T^ = MCCC 1A00836^T^ = JCM 30155^T^), isolated from seawater that was collected from the surface of the South China Sea (118° 23’ E 21° 03’ N).

### Description of *Pseudorhizobium banfieldiae* sp. nov

See protologue (Table 3) generated on the Digital Protologue Database [38] under taxon number TA00814.

**Table 3.**
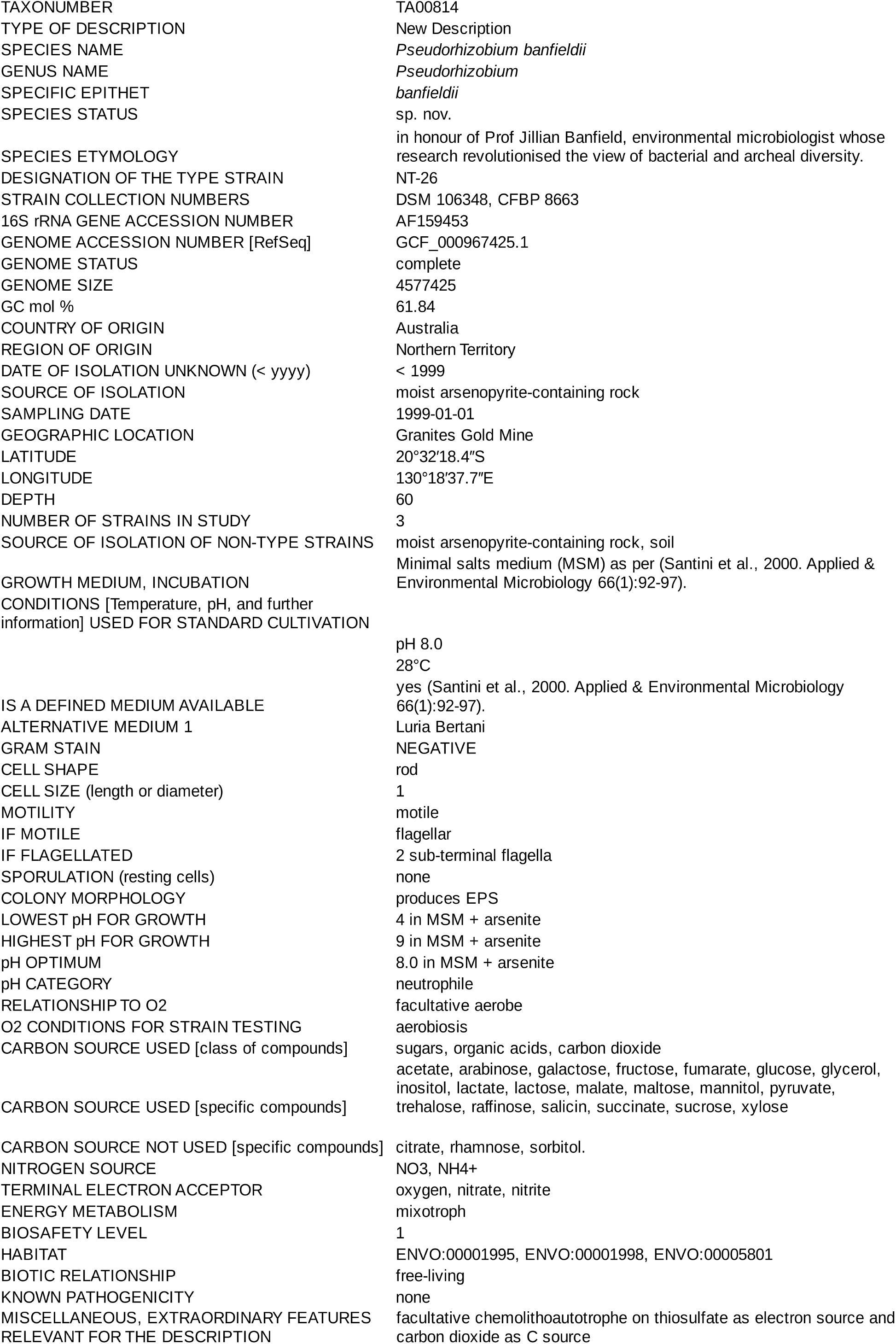
Digital Protologue description *Pseudorhizobium banfieldiae* sp. nov. (TA00814)

Etymology: N.L. gen. n. *banfieldiae* [ˈbæn · ˈfil · di · æ], named in honour of Prof Jillian Banfield, environmental microbiologist whose research revolutionised the view of bacterial and archeal diversity.

*P. banfieldiae* strains are salt tolerant up to 4% NaCl. The following phenotypes distinguish them from other members of *Pseudorhizobium*: they are sulphur oxidizers, and can harvest electrons from sulphur compounds including thiosulphate; they are autotrophic and can assimilate carbon from CO_2_ in the presence of an electron source, such as the reduced inorganic sulphur compound thiosulphate. Note that arsenite oxidation and autotrophy in the presence of arsenite are accessory traits borne by a plasmid and are not diagnostic of the species.

In addition, growth of *P. banfieldiae* strains is inhibited on yeast extract – mannitol agar medium (YEM) supplemented with 0.1g/l Congo Red dye, resulting on small, dry colonies, and are coloured orange-red by the dye.

The type strain is NT-26^T^ (= DSM 106348^T^ = CFBP 8663^T^), isolated from arsenopyrite-containing rock in a sub-surface goldmine in the Northern Territory, Australia.

## Data availability

All genomic data were submitted to the EBI-ENA under the BioProject accession PRJEB21840/ERP024139, in relation to BioSample accessions ERS1921026–ERS1921030. PacBio runs were submitted under the experiment accessions ERX2989729–ERX2989733. Illumina runs were submitted under the experiment accession ERX3427879, ERX3427880, ERX3427882– ERX3427884, ERX3431115 and ERX3431116. Annotated assemblies were submitted under the analysis accessions ERZ1027023–ERZ1027030.

Intermediary data and results from phenotypic and evolutionary analyses are available on the Figshare data repository under project 65498, available at: https://figshare.com/projects/Taxonomy_of_the_bacterial_genus_Pseudorhizobium/65498. It contains file sets relating to:

- the core genome gene alignment concatenate and the species trees *S*_ML571_, *S*_BA41_ and *T*_BA41_ (doi: 10.6084/m9.figshare.8316827);
- individual marker gene and MLSA phylogenies (doi: 10.6084/m9.figshare.8332706);
- the pangenome gene alignments for the ‘571Rhizob’ and ‘41NeoPseudo’ datasets (doi: 10.6084/m9.figshare.8343473 and doi: 10.6084/m9.figshare.8335265)
- the pangenome gene trees for the ‘41NeoPseudo’ dataset (doi: 10.6084/m9.figshare.8320199);
- the *Pantagruel* phylogenomic database summarizing the pangenome analysis of the ‘571Rhizob’ datasets (with focus on the included ‘41NeoPseudo’ dataset) (doi: 10.6084/m9.figshare.8320142);
- the fatty acid profiling of *Pseudorhizobium* strains (doi: 10.6084/m9.figshare.8316383.v1);
- the API 20 NE biochemical profiling of *Pseudorhizobium* strains (doi: 10.6084/m9.figshare.8316770);
- the NaCl Plate phenotype of *Pseudorhizobium* strains (doi: 10.6084/m9.figshare.8316803);
- the Biolog Gen III metabolism profiling of *Pseudorhizobium* strains (doi: 10.6084/m9.figshare.8316746);
- the genome-wide association testing of Biolog GenIII phenotypes vs. accessory genome presence/absence (doi: 10.6084/m9.figshare.8316818).

## Authors Contribution

FL, JMS, JP and FB designed the study. SMMD, FJZ, JZ, JMS, THO and JP isolated, provided and cultivated the bacterial strains. SV and AF performed phenotypic analyses. JS and FL analysed the phenotypic data. FL and HB conducted genome assemblies. FL and XD wrote the phylogenetic analysis software. FL conducted the bioinformatic and evolutionary analyses. FL, JMS, JP, FB and XD wrote the manuscript. All authors read and approve the content of the manuscript.

## Acknowledgement

We would like to thank Pascal Bartling, Brian Tindall, Sabine Gronow, Uli Nübel, Gabi Pötter, Peter Schumann and Philippe de Lajudie for bioinformatic, taxonomic and analytic support as well as very helpful discussions.

## References

[1] Andres, J., Arsène-Ploetze, F., Barbe, V., Brochier-Armanet, C., Cleiss-Arnold, J., Coppée, J.Y., Dillies, M.A., Geist, L., Joublin, A., Koechler, S., Lassalle, F., Marchal, M., Médigue, C., Muller, D., Nesme, X., Plewniak, F., Proux, C., Ramirez-Bahena, M.H., Schenowitz, C., Sismeiro, O., Vallenet, D., Santini, J.M., Bertin, P.N. (2013) Life in an arsenic-containing gold mine: genome and physiology of the autotrophic arsenite-oxidizing bacterium *Rhizobium* sp. NT-26. Genome Biol. Evol. 5(5), 934–53, Doi: 10.1093/gbe/evt061.

[2] Andres, J., Bertin, P.N. (2016) The microbial genomics of arsenic. FEMS Microbiol. Rev. 40(2), 299–322, Doi: 10.1093/femsre/fuv050.

[3] Antipov, D., Hartwick, N., Shen, M., Raiko, M., Lapidus, A., Pevzner, P.A. (2016) plasmidSPAdes: assembling plasmids from whole genome sequencing data. Bioinformatics 32(22), 3380–7, Doi: 10.1093/bioinformatics/btw493.

[4] Chun, J., Oren, A., Ventosa, A., Christensen, H., Arahal, D.R., da Costa, M.S., Rooney, A.P., Yi, H., Xu, X.-W., De Meyer, S., Trujillo, M.E. (2018) Proposed minimal standards for the use of genome data for the taxonomy of prokaryotes. Int. J. Syst. Evol. Microbiol. 68(1), 461–6, Doi: 10.1099/ijsem.0.002516.

[5] Chun, J., Rainey, F.A. (2014) Integrating genomics into the taxonomy and systematics of the *Bacteria* and *Archaea*. Int. J. Syst. Evol. Microbiol. 64(2), 316–24, Doi: 10.1099/ijs.0.054171-0.

[6] Cohan, F.M. (2017) Transmission in the Origins of Bacterial Diversity, From Ecotypes to Phyla. Microbiol. Spectr. 5(5), Doi: 10.1128/microbiolspec.MTBP-0014-2016.

[7] Collins, C., Didelot, X. (2018) A phylogenetic method to perform genome-wide association studies in microbes that accounts for population structure and recombination. PLOS Comput. Biol. 14(2), e1005958, Doi: 10.1371/journal.pcbi.1005958.

[8] Connor, T.R., Loman, N.J., Thompson, S., Smith, A., Southgate, J., Poplawski, R., Bull, M.J., Richardson, E., Ismail, M., Thompson, S.E.-., Kitchen, C., Guest, M., Bakke, M., Sheppard, S.K., Pallen, M.J. (2016) CLIMB (the Cloud Infrastructure for Microbial Bioinformatics): an online resource for the medical microbiology community. Microb. Genomics 2(9), e000086, Doi: 10.1099/mgen.0.000086.

[9] Deb, C., Stackebrandt, E., Pradella, S., Saha, A., Roy, P. (2004) Phylogenetically Diverse New Sulfur Chemolithotrophs of α-Proteobacteria Isolated from Indian Soils. Curr. Microbiol. 48(6), 452–8, Doi: 10.1007/s00284-003-4250-y.

[10] Diange, E.A., Lee, S.-S. (2013) *Rhizobium halotolerans* sp. nov., Isolated from Chloroethylenes Contaminated Soil. Curr. Microbiol. 66(6), 599–605, Doi: 10.1007/s00284-013-0313-x.

[11] Doyon, J.-P., Ranwez, V., Daubin, V., Berry, V. (2011) Models, Algorithms and Programs for Phylogeny Reconciliation. Brief. Bioinform. 12(5), 392–400, Doi: 10.1093/bib/bbr045.

[12] Engelhardt, T., Sahlberg, M., Cypionka, H., Engelen, B. (2011) Induction of prophages from deep-subseafloor bacteria. Environ. Microbiol. Rep. 3(4), 459–65, Doi: 10.1111/j.1758-2229.2010.00232.x.

[13] Gu, T., Sun, L.N., Zhang, J., Sui, X.H., Li, S.P. (2014) *Rhizobium flavum* sp. nov., a triazophos-degrading bacterium isolated from soil under the long-term application of triazophos. Int. J. Syst. Evol. Microbiol. 64(6), 2017–22, Doi: 10.1099/ijs.0.061523-0.

[14] Hahnke, S., Tindall, B.J., Schumann, P., Sperling, M., Brinkhoff, T., Simon, M. (2012) *Planktotalea frisia* gen. nov., sp. nov., isolated from the southern North Sea. Int. J. Syst. Evol. Microbiol. 62(7), 1619–24, Doi: 10.1099/ijs.0.033563-0.

[15] Haryono, M., Tsai, Y.-M., Lin, C.-T., Huang, F.-C., Ye, Y.-C., Deng, W.-L., Hwang, H.-H., Kuo, C.-H. (2018) Presence of an *Agrobacterium*-Type Tumor-Inducing Plasmid in *Neorhizobium* sp. NCHU2750 and the Link to Phytopathogenicity. Genome Biol. Evol. 10(12), 3188–95, Doi: 10.1093/gbe/evy249.

[16] Hördt, A., López, M.G., Meier-Kolthoff, J.P., Schleuning, M., Weinhold, L.-M., Tindall, B.J., Gronow, S., Kyrpides, N.C., Woyke, T., Göker, M. (2020) Analysis of 1,000+ Type-Strain Genomes Substantially Improves Taxonomic Classification of *Alphaproteobacteria*. Front. Microbiol. 11, Doi: 10.3389/fmicb.2020.00468.

[17] Kämpfer, P., Kroppenstedt, R.M. (1996) Numerical analysis of fatty acid patterns of coryneform bacteria and related taxa. Can. J. Microbiol. 42(10), 989–1005, Doi: 10.1139/m96-128.

[18] Kimes, N.E., López-Pérez, M., Flores-Félix, J.D., Ramírez-Bahena, M.-H., Igual, J.M., Peix, A., Rodriguez-Valera, F., Velázquez, E. (2015) *Pseudorhizobium pelagicum* gen. nov., sp. nov. isolated from a pelagic Mediterranean zone. Syst. Appl. Microbiol. 38(5), 293–9, Doi: 10.1016/j.syapm.2015.05.003.

[19] Kneen, B.E., Larue, T.A. (1983) Congo Red Absorption by *Rhizobium leguminosarum*. Appl. Environ. Microbiol. 45(1), 340–2.

[20] Konstantinidis, K.T., Tiedje, J.M. (2005) Towards a Genome-Based Taxonomy for Prokaryotes. J. Bacteriol. 187(18), 6258–64, Doi: 10.1128/JB.187.18.6258-6264.2005.

[21] Kumar, N., Lad, G., Giuntini, E., Kaye, M.E., Udomwong, P., Shamsani, N.J., Young, J.P.W., Bailly, X. (2015) Bacterial genospecies that are not ecologically coherent: population genomics of *Rhizobium leguminosarum*. Open Biol. 5(1), 140133, Doi: 10.1098/rsob.140133.

[22] de Lajudie, P.M., Andrews, M., Ardley, J., Eardly, B., Jumas-Bilak, E., Kuzmanović, N., Lassalle, F., Lindström, K., Mhamdi, R., Martínez-Romero, E., Moulin, L., Mousavi, S.A., Nesme, X., Peix, A., Puławska, J., Steenkamp, E., Stępkowski, T., Tian, C.-F., Vinuesa, P., Wei, G., Willems, A., Zilli, J., Young, P. (2019) Minimal standards for the description of new genera and species of rhizobia and agrobacteria. Int. J. Syst. Evol. Microbiol. 69, 1852–63, Doi: 10.1099/ijsem.0.003426.

[23] Lassalle, F., Jauneikaite, E., Veber, P., Didelot, X. (2019) Automated reconstruction of all gene histories in large bacterial pangenome datasets and search for co-evolved gene modules with Pantagruel. BioRxiv, 586495, Doi: 10.1101/586495.

[24] Lassalle, F., Muller, D., Nesme, X. (2015) Ecological speciation in bacteria: reverse ecology approaches reveal the adaptive part of bacterial cladogenesis. Res. Microbiol. 166(10), 729–41, Doi: 10.1016/j.resmic.2015.06.008.

[25] Lassalle, F., Planel, R., Penel, S., Chapulliot, D., Barbe, V., Dubost, A., Calteau, A., Vallenet, D., Mornico, D., Bigot, T., Guéguen, L., Vial, L., Muller, D., Daubin, V., Nesme, X. (2017) Ancestral Genome Estimation Reveals the History of Ecological Diversification in *Agrobacterium*. Genome Biol. Evol. 9(12), 3413–31, Doi: 10.1093/gbe/evx255.

[26] Lepage, T., Bryant, D., Philippe, H., Lartillot, N. (2007) A General Comparison of Relaxed Molecular Clock Models. Mol. Biol. Evol. 24(12), 2669–80, Doi: 10.1093/molbev/msm193.

[27] Li, H. (2016) Minimap and miniasm: fast mapping and de novo assembly for noisy long sequences. Bioinformatics 32(14), 2103–10, Doi: 10.1093/bioinformatics/btw152.

[28] Linström, K. (1989) *Rhizobium galegae*, a New Species of Legume Root Nodule Bacteria. Int. J. Syst. Evol. Microbiol. 39(3), 365–7, Doi: 10.1099/00207713-39-3-365.

[29] Liu, Y., Wang, R.-P., Ren, C., Lai, Q.-L., Zeng, R.-Y. (2015) *Rhizobium marinum* sp. nov., a malachite-green-tolerant bacterium isolated from seawater. Int. J. Syst. Evol. Microbiol. 65(12), 4449–54, Doi: 10.1099/ijsem.0.000593.

[30] Meier-Kolthoff, J.P., Auch, A.F., Klenk, H.-P., Göker, M. (2013) Genome sequence-based species delimitation with confidence intervals and improved distance functions. BMC Bioinformatics 14(1), 60, Doi: 10.1186/1471-2105-14-60.

[31] Mousavi, S.A., Österman, J., Wahlberg, N., Nesme, X., Lavire, C., Vial, L., Paulin, L., de Lajudie, P., Lindström, K. (2014) Phylogeny of the *Rhizobium*–*Allorhizobium*–*Agrobacterium* clade supports the delineation of *Neorhizobium* gen. nov. Syst. Appl. Microbiol. 37(3), 208–15, Doi: 10.1016/j.syapm.2013.12.007.

[32] Mousavi, S.A., Willems, A., Nesme, X., de Lajudie, P., Lindström, K. (2015) Revised phylogeny of *Rhizobiaceae*: Proposal of the delineation of *Pararhizobium* gen. nov., and 13 new species combinations. Syst. Appl. Microbiol. 38(2), 84–90, Doi: 10.1016/j.syapm.2014.12.003.

[33] Oren, A., Garrity, G.M. (2017) List of new names and new combinations previously effectively, but not validly, published. Int. J. Syst. Evol. Microbiol. 67(9), 3140–3, Doi: 10.1099/ijsem.0.002278.

[34] Ormeño-Orrillo, E., Servín-Garcidueñas, L.E., Rogel, M.A., González, V., Peralta, H., Mora, J., Martínez-Romero, J., Martínez-Romero, E. (2015) Taxonomy of rhizobia and agrobacteria from the *Rhizobiaceae* family in light of genomics. Syst. Appl. Microbiol. 38(4), 287–91, Doi: 10.1016/j.syapm.2014.12.002.

[35] Österman, J., Marsh, J., Laine, P.K., Zeng, Z., Alatalo, E., Sullivan, J.T., Young, J.P.W., Thomas-Oates, J., Paulin, L., Lindström, K. (2014) Genome sequencing of two *Neorhizobium galegae* strains reveals a noeT gene responsible for the unusual acetylation of the nodulation factors. BMC Genomics 15(1), 500, Doi: 10.1186/1471-2164-15-500.

[36] Parag, B., Sasikala, C., Ramana, C.V. (2013) Molecular and culture dependent characterization of endolithic bacteria in two beach sand samples and description of *Rhizobium endolithicum* sp. nov. Antonie Van Leeuwenhoek 104(6), 1235–44, Doi: 10.1007/s10482-013-0046-7.

[37] Parks, D. (2020) dparks1134/CompareM.

[38] Rosselló-Móra, R., Trujillo, M.E., Sutcliffe, I.C. (2017) Introducing a digital protologue: a timely move towards a database-driven systematics of archaea and bacteria. Antonie Van Leeuwenhoek 110(4), 455–6, Doi: 10.1007/s10482-017-0841-7.

[39] Santini, J.M., vanden Hoven, R.N. (2004) Molybdenum-Containing Arsenite Oxidase of the Chemolithoautotrophic Arsenite Oxidizer NT-26. J. Bacteriol. 186(6), 1614–9, Doi: 10.1128/JB.186.6.1614-1619.2004.

[40] Santini, J.M., Sly, L.I., Schnagl, R.D., Macy, J.M. (2000) A New Chemolithoautotrophic Arsenite-Oxidizing Bacterium Isolated from a Gold Mine: Phylogenetic, Physiological, and Preliminary Biochemical Studies. Appl. Environ. Microbiol. 66(1), 92–7, Doi: 10.1128/AEM.66.1.92-97.2000.

[41] Schumann, P., Maier, T. (2014) Chapter 13 - MALDI-TOF Mass Spectrometry Applied to Classification and Identification of Bacteria. In: Goodfellow, M., Sutcliffe, I., Chun, J., (Eds.), Methods in Microbiology, vol. 41, Academic Press, pp. 275–306.

[42] Simon, M., Scheuner, C., Meier-Kolthoff, J.P., Brinkhoff, T., Wagner-Döbler, I., Ulbrich, M., Klenk, H.-P., Schomburg, D., Petersen, J., Göker, M. (2017) Phylogenomics of *Rhodobacteraceae* reveals evolutionary adaptation to marine and non-marine habitats. ISME J. 11(6), 1483–99, Doi: 10.1038/ismej.2016.198.

[43] Stackebrandt, E., Frederiksen, W., Garrity, G.M., Grimont, P.A.D., Kämpfer, P., Maiden, M.C.J., Nesme, X., Rosselló-Mora, R., Swings, J., Trüper, H.G., Vauterin, L., Ward, A.C., Whitman, W.B. (2002) Report of the ad hoc committee for the re-evaluation of the species definition in bacteriology. Int. J. Syst. Evol. Microbiol. 52(Pt 3), 1043–7.

[44] Surange, S., Wollum II, A.G., Kumar, N., Nautiyal, C.S. (1997) Characterization of *Rhizobium* from root nodules of leguminous trees growing in alkaline soils. Can. J. Microbiol. 43(9), 891–4, Doi: 10.1139/m97-130.

[45] Suzuki, R., Shimodaira, H. (2006) Pvclust: an R package for assessing the uncertainty in hierarchical clustering. Bioinformatics 22(12), 1540–2, Doi: 10.1093/bioinformatics/btl117.

[46] Suzuki, T., Campbell, J., Kim, Y., Swoboda, J.G., Mylonakis, E., Walker, S., Gilmore, M.S. (2012) Wall teichoic acid protects *Staphylococcus aureus* from inhibition by Congo red and other dyes. J. Antimicrob. Chemother. 67(9), 2143–51, Doi: 10.1093/jac/dks184.

[47] Szöllősi, G.J., Rosikiewicz, W., Boussau, B., Tannier, E., Daubin, V. (2013) Efficient Exploration of the Space of Reconciled Gene Trees. Syst. Biol. 62(6), 901–12, Doi: 10.1093/sysbio/syt054.

[48] Teufel, R., Mascaraque, V., Ismail, W., Voss, M., Perera, J., Eisenreich, W., Haehnel, W., Fuchs, G. (2010) Bacterial phenylalanine and phenylacetate catabolic pathway revealed. Proc. Natl. Acad. Sci. 107(32), 14390–5, Doi: 10.1073/pnas.1005399107.

[49] Tóth, E.M., Schumann, P., Borsodi, A.K., Kéki, Z., Kovács, A.L., Márialigeti, K. (2008) *Wohlfahrtiimonas chitiniclastica* gen. nov., sp. nov., a new gammaproteobacterium isolated from *Wohlfahrtia magnifica* (*Diptera*: *Sarcophagidae*). Int. J. Syst. Evol. Microbiol. 58(4), 976–81, Doi: 10.1099/ijs.0.65324-0.

[50] Vaas, L.A.I., Sikorski, J., Hofner, B., Fiebig, A., Buddruhs, N., Klenk, H.-P., Göker, M. (2013) opm: an R package for analysing OmniLog(R) phenotype microarray data. Bioinforma. Oxf. Engl. 29(14), 1823–4, Doi: 10.1093/bioinformatics/btt291.

[51] Wang, E.T., van Berkum, P., Beyene, D., Sui, X.H., Dorado, O., Chen, W.X., Martínez-Romero, E. (1998) Rhizobium huautlense sp. nov., a symbiont of *Sesbania herbacea* that has a close phylogenetic relationship with *Rhizobium galegae*. Int. J. Syst. Evol. Microbiol. 48(3), 687–99, Doi: 10.1099/00207713-48-3-687.

