## Supplementary material for "Phylogenomic analysis reveals the basis of adaptation of *Pseudorhizobium* species to extreme environments": Sup. Fig.

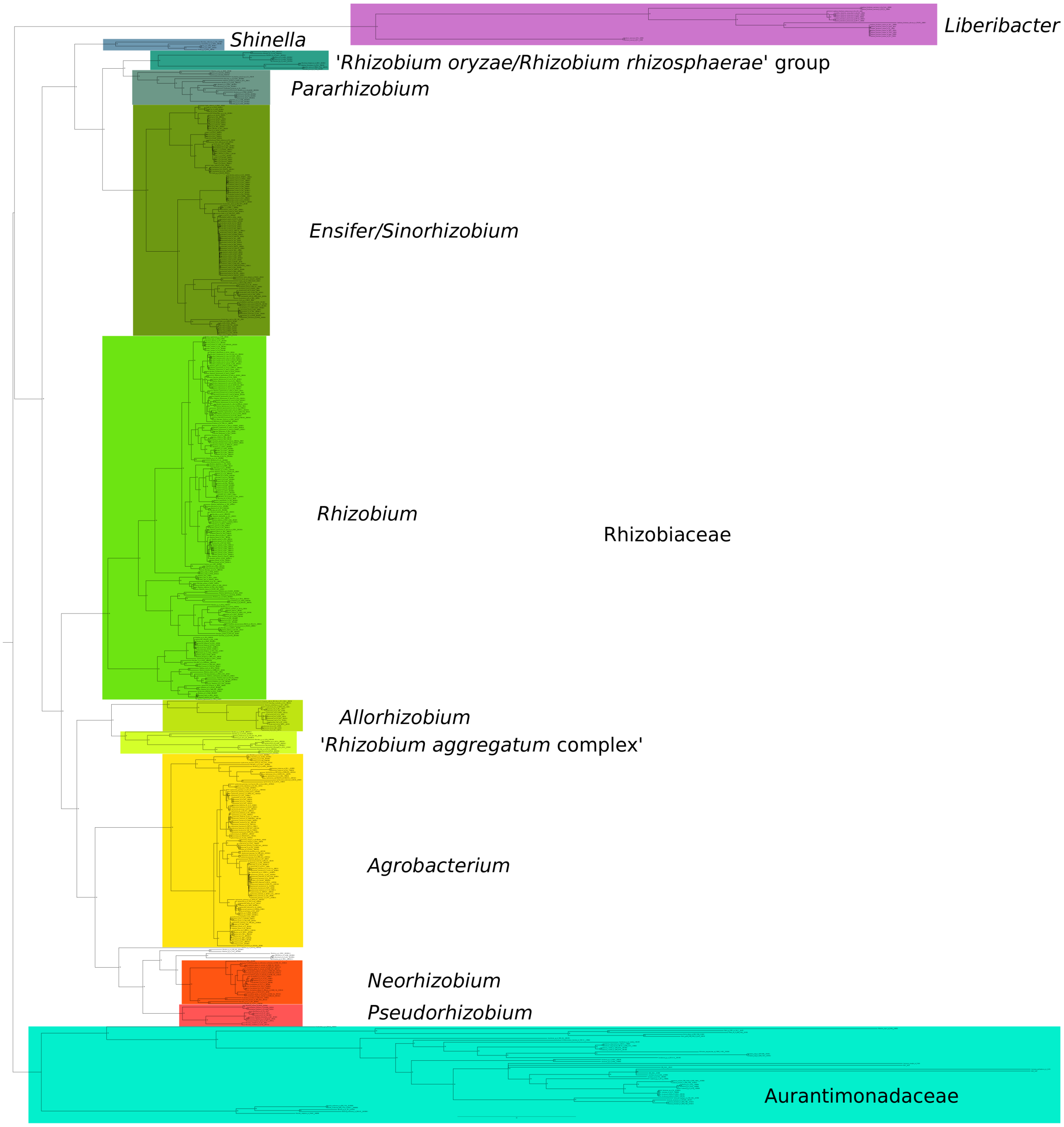


**Figure S1. Maximum-likelihood core genome phylogeny of 571 *Rhizobiaceae* and *Aurantimonadaceae* ($S_{ML571}$).**
Tree was obtained with RAxML under the model PROTCATLGX based on the concatenated alignment of 155 core protein families occurring only in a single copy and present in at least 561 out of the 571 genomes (98%); branch supports were estimated by generating 200 rapid bootstraps under the same parameters. The alignment and tree files are available on Figshare (doi: 10.6084/m9.figshare.8316827.v1).

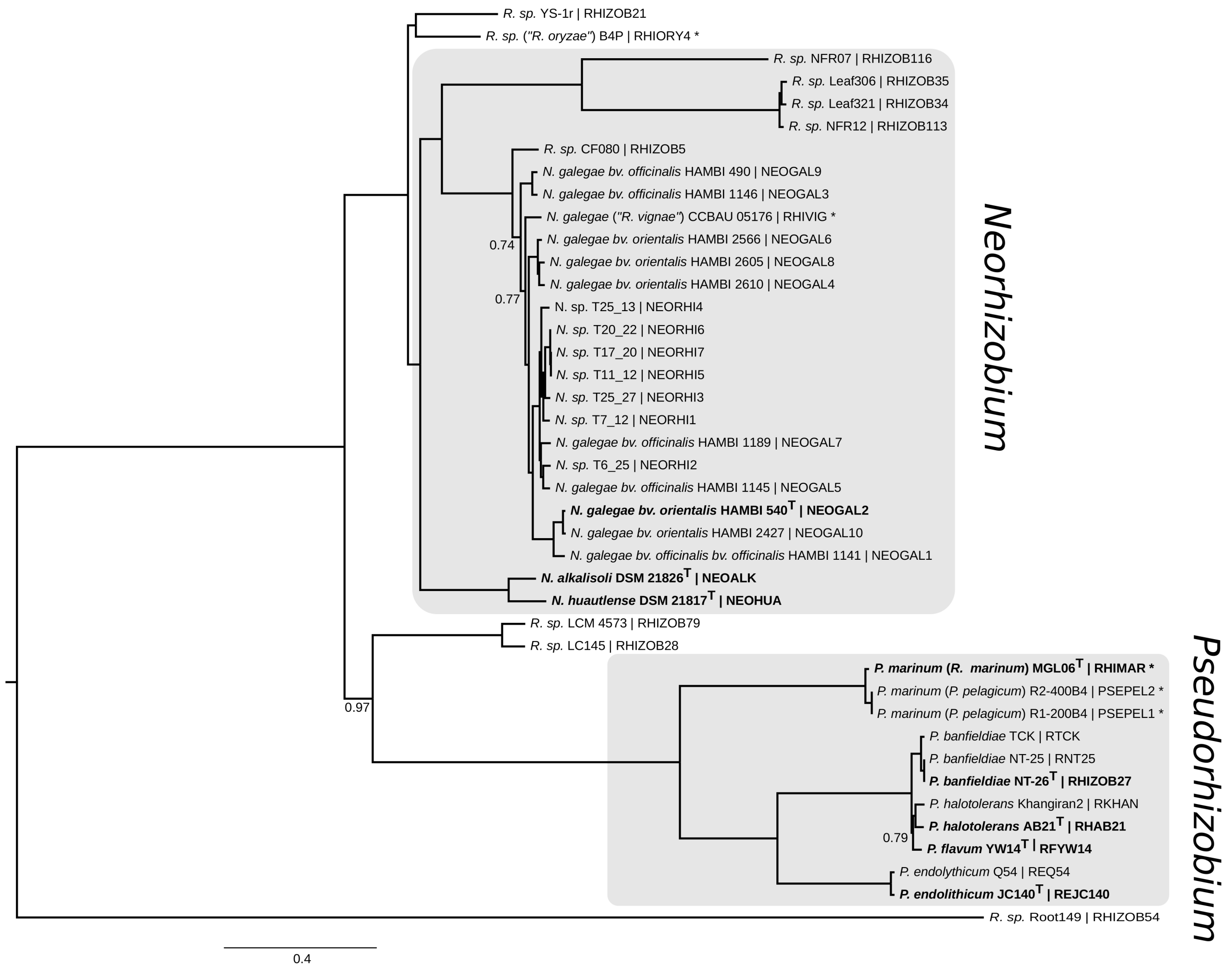


**Figure S2. Bayesian phylogenetic tree of 41 organisms from the *Neorhizobium* and *Pseudorhizobium* genera and close relatives ($S_{BA41}$).**

Same tree as in Figure 1, presented in full. Tree obtained with Phylobayes under the GTR-CAT protein evolution model, based on a concatenated alignment of 155 pseudo-core protein loci. All posterior probability branch supports are 1.00 unless indicated. The organism name is followed (after the pipe symbol | ) by the identifier in the Pantagruel pangenome database of this study. Strains whose species affiliation are corrected in this study are marked with an asterisk *. Species type strains are in bold. The alignment and tree files are available on Figshare (doi: 10.6084/m9.figshare.8316827.v1).

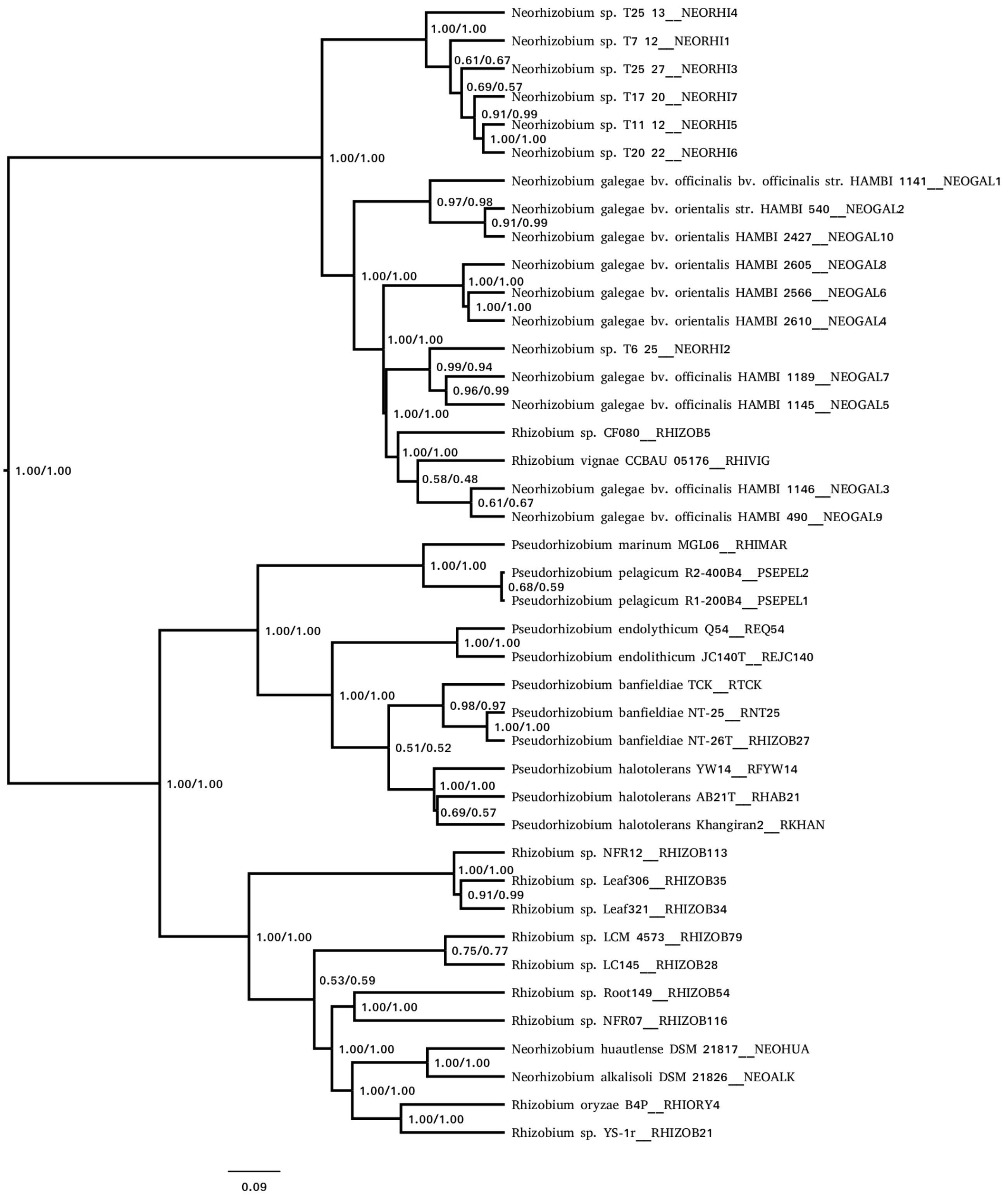


**Figure S3. Hierarchical clustering of 41 organisms from the *Neorhizobium* and *Pseudorhizobium* genera and close relatives ($S_{CL41}$).**

Dendrogram obtained by the Ward (D2) clustering method based on the Jaccard distance between genomes derived from the distribution of orthologous groups of accessory genes. Numbers at nodes indicate supports from bootstraping and approximately unbiased (AU) test, as obtained with pvclust R package.

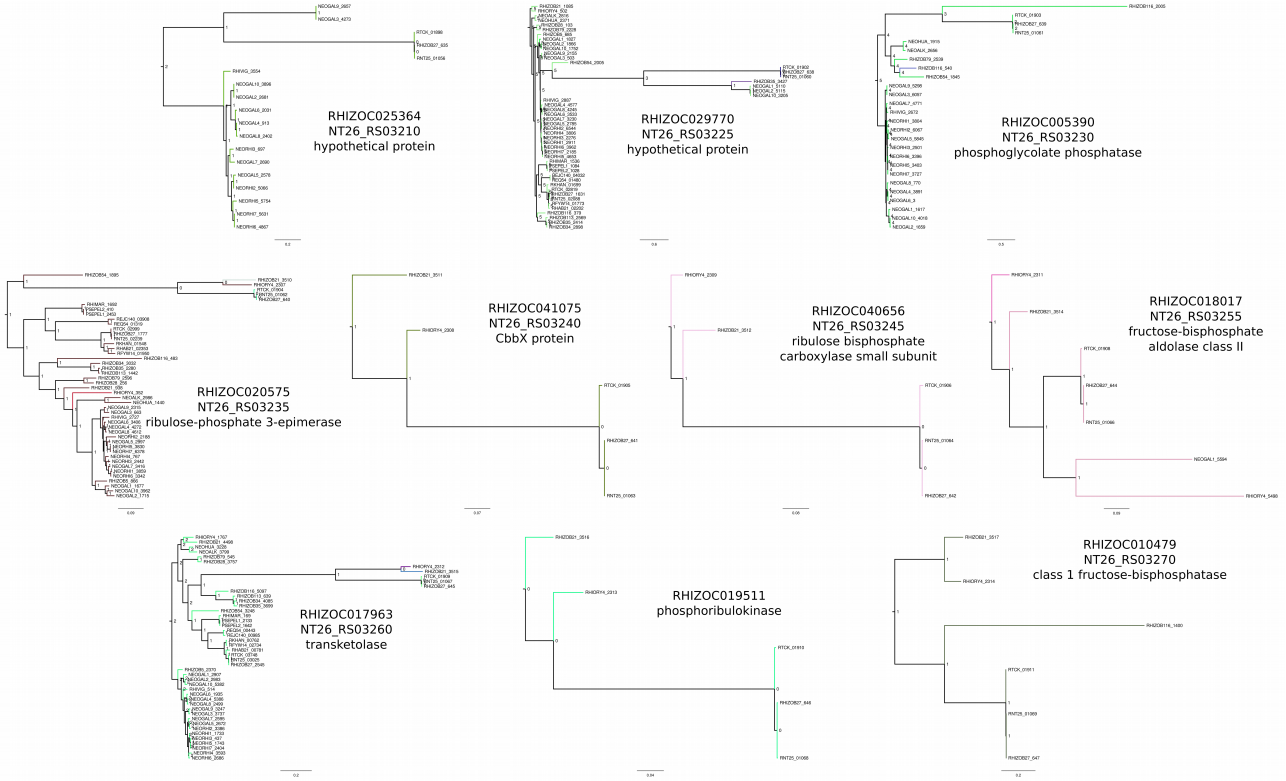


**Fig. S4. Reconciled gene trees of the gene families composing the RuBisCO operon in *Pban*.**

For each gene family is indicated the family id, the gene locus tag in NT-26 genome, and its functional annotation. In each tree, tips are coloured by orthologous group (OG); a change of OG between a clade and the clade from which it emerges indicates a gene gain event (see Sup. Table S4 for event scenarios including the list of HGT events). Gene trees and event records are available on Figshare (doi: 10.6084/m9.figshare.8320199.v1 and doi: 10.6084/m9.figshare.8320142.v1).

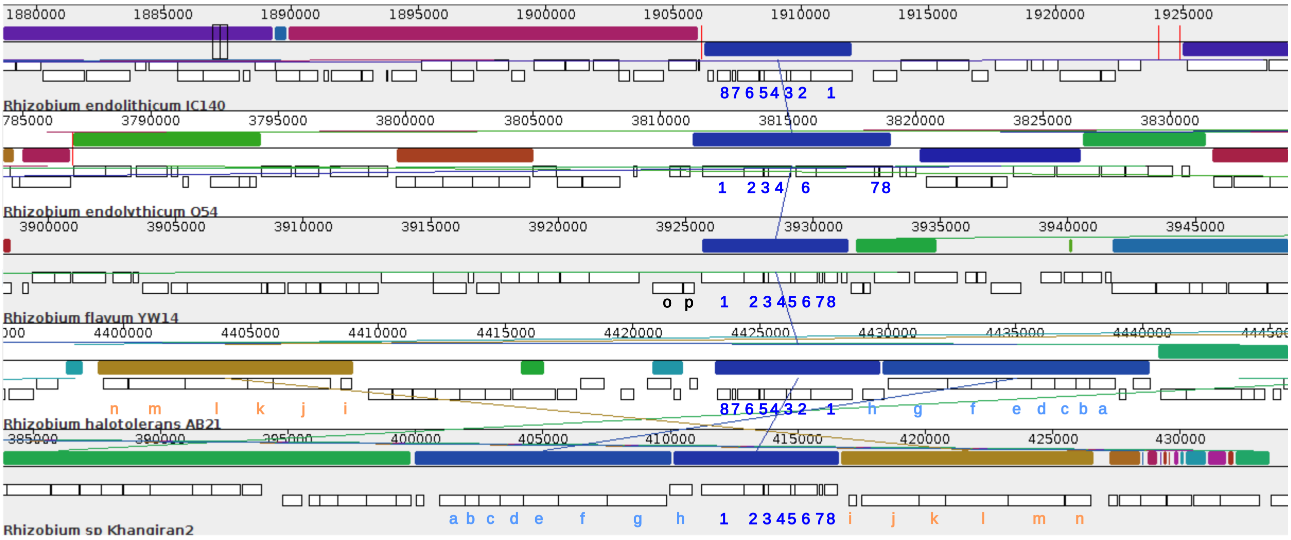


**Figure S5. Syntenic map of cytochrome-c oxidase and *eutK* locus in *Pfla*, *Phalo* and *Pendo* genomes.**

The map is a snapshot from Mauve viewer of a multiple genome alignment generated with progressiveMauve. Homologous genome segments, or local collinear blocks (LCBs), are coloured the same and linked with an edge. White boxes indicate annotated coding sequences. Letters and digit refer to the functional annotation below: 1-8: cytochrome-c oxidase and *eutK* locus; a-h: nitrous oxide reductase locus; i-n: cellulose synthase locus; o-p: nitrite reductase locus.

1: cytochrome-c oxidase, cbb3-type subunit I
2: cytochrome-c oxidase, cbb3-type subunit II
3: CcoQ/FixQ family Cbb3-type cytochrome c oxidase assembly chaperone
4: cytochrome-c oxidase, cbb3-type subunit III
5: hypothetical protein
6: hypothetical protein
7: cbb3-type cytochrome oxidase assembly protein CcoS
8: ethanolamine utilization microcompartment protein EutK
a: FAD:protein FMN transferase
b: NosL copper chaperone
c: membrane protein
d: ABC transporter ATP-binding protein
e: nitrous oxide reductase family maturation protein NosD
f: TAT-dependent nitrous-oxide reductase
g: regulatory protein NosR
h: rubrerythrin family protein
i: hypothetical protein
j: cellulose synthase
k: endoglucanase
l: cellulose synthase BcsB subunit
m: cellulose synthase catalytic subunit (UDP-forming)
n: cellulose biosynthesis protein BcsN
o: nitrite reductase, copper-containing
p: pseudoazurin

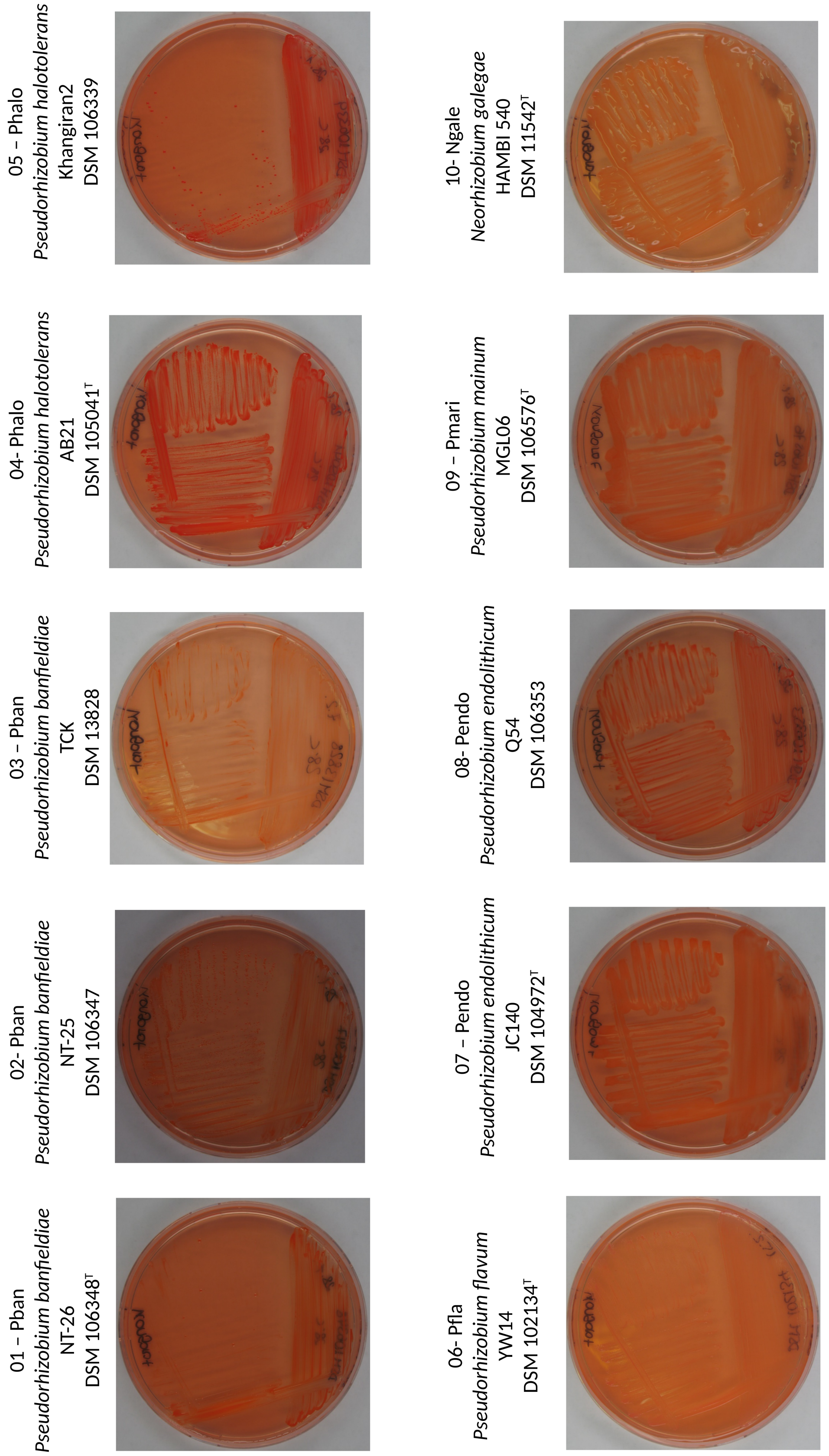


**Figure S6. Congo Red assay.**

Strains were plated on yeast mannitol agar medium (YEM) with 0.1 g/L Congo red dye for seven days. YEM composition: 10g mannitol, 1g yeast extract, 9.5 $K_2HPO_4$, 0.2g $MgSO_4$, $7H_2O$, 0.1g NaCl (Surrange et al., 1998. *Can J Microbiol* 43:891-894). See Sup. Table S7 for classification of strain growth patterns.

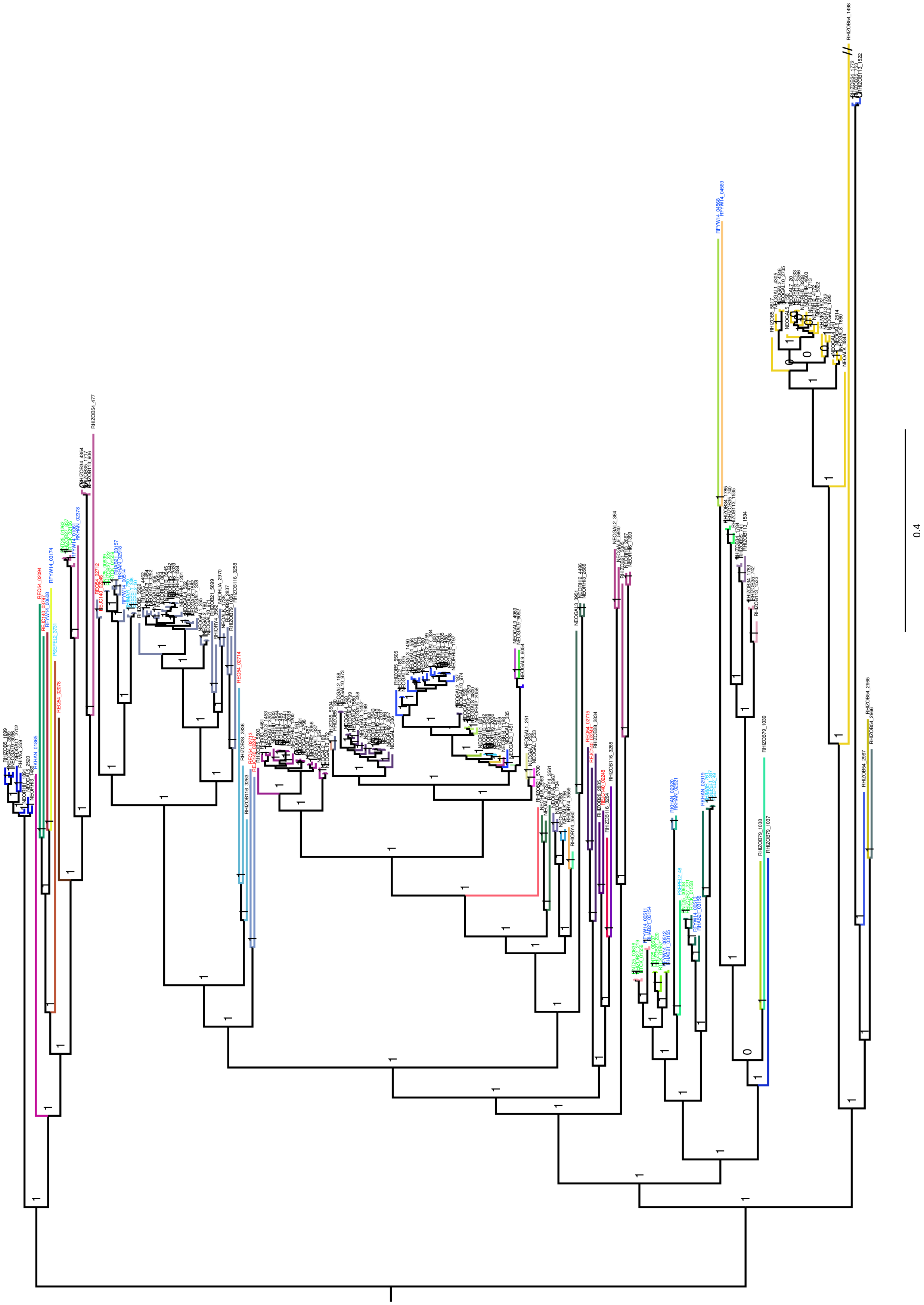

**Figure S7. Phylogenetic tree of flagellin gene family (RHIZOC021332).**

Tips are coloured by group of orthologs within this large multi-copy gene family (as inferred based on the unicity criterion). Sequence belonging to relevant organisms have their labels coloured in red (*Pendo*), dark blue (*Phalo+Pfla*), light blue (*Pmari*) or green (*Pban*).
