## Supplementary material for "Phylogenomic analysis reveals the basis of adaptation of *Pseudorhizobium* species to extreme environments": Suppl. Text

### Supplementary Methods

#### Genomic DNA quality control

For the strains *Rhizobium* sp. NT-25, *R. flavum* YW14, *R.* sp. Q54, *R.* sp. TCK and Khangiran2, quality of the DNA (notably fragment size) was checked by electrophoresis on 0.5% agarose gel and the identity and purity of the DNA was checked by amplifying the *recA* marker gene by PCR using *Rhizobiaceae*-specific primers F7386 and F7387 (Shams et al. 2013), Sanger sequencing and comparison to known sequences (NCBI GenBank accessions for partial *recA* gene sequences KF609368.1, KF609367.1, JX872399.1 and extracted CDS of gene NT26_RS09070 from chromosome sequence NZ_FO082820.1). Genomic DNA was sent to the Earlham Institute for sequencing, where quantity and quality of the DNA was determined using a Qubit fluorometer (Invitrogen, Waltham, Massachusetts) and fragment length was controlled with Tapestation 2200 (Agilent, Santa Clara, California).

#### Phylogenomic analyses

Unless specified otherwise, the following bioinformatic analyses were conducted using the Pantagruel under default setting as described in (Lassalle et al. 2019) and on the program webpage http://github.com/flass/pantagruel/. This pipeline is designed for the analysis of bacterial pangenomes, inluding the inference of a species tree, gene trees, and the detection of horizontal gene transfers through specie tree/gene tree reconciliations (Szöllősi et al. 2015)
.

##### *Genome annotation*

*Pantagruel task 0*: we used Prokka (version 1.12) (Seemann 2014) to annotate the new genomes sequences, with the following notable options: “--force --addgenes --locustag --compliant –usegenus –genus Rhizobium --kingdom Bacteria --gcode 11”. We used a reference database of annotated protein based on proteomes predicted from all available complete genomes from the *Rhizobium*/*Agrobacterium* group, as downloaded from the NCBI RefSeq Assembly database on the 15 Dec 2017 using the query ‘txid227290[Organism:exp] AND ("latest refseq"[filter] AND ("scaffold level"[filter] OR "chromosome level"[filter] OR "complete genome"[filter]) AND all[filter] NOT anomalous[filter])’. These proteomes were merged into a non-redundant dataset that was thinned by retaining a single representative protein for each protein cluster as determined by the CD-HIT clustering program (version 4.6) (Fu et al. 2012), used with the options ‘-T 0 -M 0 -G 1 -s 0.8 -c 0.9’. The resulting dataset was used as a subject database for a BLASTP (Camacho et al. 2009) similarity search during the annotation process.

##### *Genomic dataset and gene family classification*

We assembled a complete bacterial genome dataset covering the known diversity subgroup of the alphaproteobacterial families *Rhizobiaceae* and (sister group) *Aurantimonadaceae*. This dataset comprises all 564 genomes available from the NCBI RefSeq Assembly database on the 23 Apr 2018, filtering anomalous genomes and those with a contig N50 < 98kb using query: ‘(txid82115[Organism:exp] OR txid255475[Organism:exp]) AND ("latest refseq"[filter] NOT anomalous[filter]) AND ("98000"[ContigN50] : "20000000"[ContigN50])’. From this dataset we removed assembly GCF_002000045.1 (*R. flavum* YW14) for which we had a newer, higher quality assembly. To these we added the new genome sequences from the eight strains mentioned above, for a total of 571 genomes (dataset ‘571Rhizob’).

*Pantagruel task 1*: all predicted protein sequence were extracted from the dataset of 571 annotated genomes and clustered into homologous gene families (with possibly multiple copies per genome) using MMSeqs2 release 2 (commit e5d64b2) (Steinegger & Söding 2017)
.

*Pantagruel task 2*: Protein sequences from each family were aligned using ClustalOmega (Sievers et al. 2011) and reverse-translated into coding sequence (CDS) alignments using PAL2NAL (Suyama et al. 2006).

*Pantagruel task 3*: a relational database (with a SQLite engine) was created to gather all annotation data as well as homologous gene classification, and data from further steps.

*Pantagruel task 4*: in addition to Prokka annotation, functional annotation of proteins was done using the standalone InterProScan tool (version 5.32-71.0) (Jones et al. 2014) with options ‘ --iprlookup --goterms –pathways’ to add annotation of InterPro terms, Gene Ontology (GO) terms and KEGG, BioCyc and Reactome metabolic pathways, providing an annotation with controlled vocabulary that is homogeneous across the dataset.

*Reference species trees*

*Pantagruel task 5*: We used this dataset of 571 annotated genomes to define the pseudo-core genome as genes occurring only in a single copy and present in at least 561 out of the 571 genomes (98%). The resulting pseudo-core genome gene set (thereafter referred as $pCG_{571}$) includes 155 loci, which protein alignments were concatenated. This concatenate alignment was used to compute a reference species tree ($S_{ML571}$) with RAxML version 8.2.11 (Stamatakis 2014) under the model PROTCATLGX; branch supports were estimated by generating 200 rapid bootstraps under the same parameters.

(Extra analyses not part of *Pantagruel* workflow:) From the $S_{571}$ tree, we identified the well-supported clade grouping 41 genomes including all representative of *Neorhizobium* spp. and *Pseudorhizobium* spp. and our new isolates (dataset ‘41NeoPseudo’). To gain further phylogenetic resolution in this clade of interest, we restricted the $pCG_{571}$ concatenated alignment to the 41 genomes of this smaller genomic dataset, which we used as input for a more accurate (but computationally more expensive) Bayesian phylogenetic inference. We used the program Phylobayes-MPI program version 1.8 to estimate a reference species phylogeny under the CAT-GTR amino-acid equilibrium frequency profile mixture model (Lartillot et al. 2007) with four site categories of discrete Gamma distribution of relative substitution rates (options ‘-cat -gtr -dgam 4’). We ran two independent chains until convergence of the topology (bipartition convergence diagnostics on 5,196 generations, excluding the first 1,500 as burn-in sample: maxdiff = 0.25; meandiff < 0.01); this provided us with a robust non-ultrametric tree for the 41 genomes ($S_{BA41}$). We finally used this $S_{BA41}$ tree as a fixed input topology for Phylobayes version 4.1c under the CIR clock model (Lepage et al. 2007), for which we ran two independent chains until convergence of the rate parameters to estimate an ultrametric (unitless ‘time’) tree ($T_{BA41}$).

*Gene trees, reconciliations and orthologous group classification*

*Pantagruel task 6*: gene trees were computed for each of the 6,714 homologous gene family of the 41-species pangenome with at least 4 sequences using MrBayes version 3.2.6 (Ronquist & Huelsenbeck 2003) under the GTR+4G+I model (with command 'lset nst=6 rates=invgamma ploidy=haploid'), running the Metropolis-coupled Markov chain Monte-Carlo (MCMCMC) for 2,000,000 generations with four parallel chains (one cold and three gradually heated) in each of two independent runs, sampling a tree every 500 generations. Convergence of bipartition distribution in pairs of 4,000 sampled gene tree sets was achieved for all gene families under these conditions.

*Pantagruel task 7*: dated gene tree/species tree reconciliation was conducted for each gene family to infer their scenarios of evolution, with events of gene duplication, transfer and loss (DTL) using ALE version 0.4 (Szöllősi, Tannier, et al. 2013)
(Szöllősi, Tannier, et al. 2013). We combine the pair of gene tree samples (resulting in 8,000 trees/gene family) and used them as input for the ALEml program, discarding the first 2,000 trees as burn-in, to be reconciled with the dated tree $T_{BA41}$. Gene DTL event rate parameters were set free and estimated by a maximum-likelihood (ML) approach, and then 1,000 gene family evolution scenarios and associated reconciliations were sampled using a Bayesian approach (Szöllősi, Rosikiewicz, et al. 2013)
(Szöllősi, Rosikiewicz, et al. 2013)

*Pantagruel task 8*: based on the estimated gene family evolution scenarios, we could define gene groups based on a true criterion of orthology, i.e. common descent from an ancestor by means of speciation only (Doyon et al. 2011), rather than a proxy criterion such as bidirectional best hits (BBH) in a similarity search. This has the advantage of explicitly detecting the gain of an orthologous gene group (OGG) in a genome lineage by means of horizontal gene transfer (HGT) or gene duplication. To differentiate additive HGT and replacing HGT events (i.e. gene conversion by homologous recombination events), we used a heuristic based on the unicopy criterion (Bigot et al. 2013) as described in a previous study (Lassalle et al. 2017), where transfer events that do not increase the gene copy number in a genome are not interpreted as the emergence of a new OGG, i.e. considering homologous recombination can occur within an OGG without breaking it. This OGG classification heuristic was applied to each sampled scenario among the 1,000 trees sampled for each gene family. To summarize these data, a network was built that connects genes classified in the same OGG in more than 50% of the sample; connected components of this graph provided the final consensus OGGs. We then used this classification to build a matrix of OGG presence/absence in the '41NeoPseudo' dataset, and computed the clades-specific core genome gene set for each clade of the species tree $S_{BA41}$, under the following criteria: 1) every OGG present in all members of the focal clade and absent in all members of its sister clade, and 2) for chosen clades matching *Pseudorhizobium* constituent species, every OGG present in all members of the species, and absent in the rest of the *Pseudorhizobium* genus.

Hierarchical clustering was performed based on the OGG presence/absence matrix using the pvclust function form the pvclust R package version 2.0-0 (Suzuki & Shimodaira 2006) with default settings, to obtain bootstrap-derived p-values (BP) and approximately unbiased (AU) branch support estimates.

*Gene Ontology enrichment tests*

GO terms frequency of annotation (as derived from the automatic InterProScan annotation) were compared between test ('study') gene sets and respective reference ('population') gene sets, where the former is included in the latter. Gene set pairs that were compared comprised: 1) clade-specific core genome of a clade (a single representative sequence per OGG) vs. whole core genome of this clade (a single representative sequence per OGG); and 2) clade-specific core genome of a clade (all sequences) vs. pangenome of this clade (all sequences). All tests were performed using the topGO R package version 2.22.0 (Alexa & Rahnenfuhrer 2010), using the 'weight01' algorithm and 'Fisher' statistic.

*Phenotype association testing*

The association of accessory gene occurrence with phenotypic profiles obtained with the Biolog GenIII (continuous values) was tested using the phylogenetic framework implemented in the 'treeWAS' R package (Collins & Didelot 2018). The test was performed on 9,621 non-trivial OGG presence/absence profiles using the ML method for ancestral state reconstruction. Significance was assessed by comparing test scores to those obtained by simulating random genotype profiles along the species tree (10-taxon tree derived from $S_{BA41}$) for 100 times more loci than the tested dataset (ca. $10^5$ simulations) and scaling the *p*-values using FDR correction. Phenotypes with significant hits were then tested again with more stringent parameters, using 1,000 times more loci than the tested dataset and Bonferroni *p*-value correction.

*Marker gene phylogenies*

CDS alignments and gene trees were extracted from the '41NeoPseudo' Pantagruel database described above for *atpD* (gene family id: RHIZOC034665, OGG id: 1), *glnII* (RHIZOC002590, 0), *gyrB* (RHIZOC002739, 0), *recA* (RHIZOC040013, 0) and *rpoB* (RHIZOC004093, 0). Note that *R. marinum* MGL06$^T$ sequence for *glnII* gene was missing in gene family RHIZOC002590, probably due to incomplete genome assembly. CDS alignments were concatenated and a MLSA tree was computed with MrBayes as described above for single gene trees.

### Supplementary Text

#### Alternative sources of taxonomic classification information

We first evaluated whther the distribution of pangenome genes could be used as an alternative to core genome alignements to infer the evolutionary relationships between strains. A hierarchical clustering based on the distribution of groups of orthologous accessory genes present in the '41NeoPseudo' genomes ($S_{CL41}$) shows a very similar picture to $S_{BA41}$, with good support for most branches leading to major clades and species (Figure 2B; Sup. Fig. S3). The branch separating *Pfla* YW14$^T$ from strains Khangiran2 and *Phalo* AB21$^T$ had low support (BP: 0.51, AU: 0.52) indicating that only half the accessory genes support the presented topology; the other half is distributed in a non-parsimonious way, suggestive a frequent HGT between these two species. A major difference with the core genome tree $S_{BA41}$ is that deep-branching strains in $S_{BA41}$ all cluster as a sister group of the *Pseudorhizobium* clade in the pangenome tree $S_{CL41}$ (Sup. Fig. S3). This illustrates that while gene presence/absence provides an interesting complement of data possibly allowing a better resolution at the genus level and below, its interpretation in terms of evolutionary history should be only complementary to (core genome) sequence-based data.

Finally, we compared these phylogenies based on core genome-wide and pangenome-wide data to those obtained using only a restricted set of classic marker genes, either separately or in concatenation, i.e. in a multi-locus sequence analysis (MLSA) (Sup. Dataset S1). The monophyly of *Pmari* and the monophyly of *Pendo* are consistently recovered for each of the five tested marker genes *atpD*, *recA*, *rpoB*, *glnII* and *gyrB*. However, only the *atpD*, *rpoB*, *glnII* and *gyrB* trees, but not *recA*, support *Pban* to be monophyletic. Also, none of these genes support the monophyly of the group *Phalo*+*Pfla*, specifically due to the inclusion of the *Pban* clade in the former. Combining these markers, the MLSA finds *Pmari*, *Pendo* and *Pban* as being monophyletic, but again not *Phalo*+*Pfla*, with the inclusion of *Pban* being moderately supported (PP support < 0.86). These results indicate that while marker gene-based analyses are mostly consistent with the information obtained from whole genomes, they do not allow for a systematically correct classification of strains into species.

**Clade-specific gene sets reveal specific functions and ecologies**

We inferred gene family evolution scenarios accounting for HGT history by reconciling gene tree topologies with that of the species tree $S_{BA41}$. Based on these scenarios, we delineated groups of orthologous genes that reflect the history of gene acquisition in genome lineages – a new orthologous cluster being delineated every time a new gene copy was gained in a lineage. We then searched gene orthologs that were gained or lost specifically by the clade ancestors of the $S_{BA41}$ tree and which presence/absence state was conserved thereafter by the clade members, i.e. genes which have occurrence patterns in full contrast between the clade and its sister group (referred to as 'clade-specific' genes), or between a species and all other species in the genus (stricter criterion, referred to as 'species-specific' genes). Data for all clade comparisons in our '41NeoPseudo' dataset are presented in Sup. Table S3 and are summarized below for the main clades of interest: the 'NT-25/26 clone', '*Pseudorhizobium banfieldiae*' (*Pban*), '*Pseudorhizobium halotolerans*' (*Phalo*), '*Pseudorhizobium endolithicum*' (*Pendo*) and '*P. marinum*' (*Pmari*). Major gene sets that have contrasting pattern of occurrence in *Pseudorhizobium* are listed Table 2 and those specifically contributing to the differentiation of the NT-25/26 clone lineage are depicted in Figure 3.

*'NT-25/NT-26 clone'*

Most of the genome gene content specific to this group is made of mobile or selfish elements. As expected, this includes the plasmids carrying the arsenite oxidation *aio* locus, the arsenic resistance *ars* operon and the phosphate-specific transporter *pst*/*phn* locus, found on a 322-kb plasmid in NT-26 and a 119-kb plasmid in NT-25. In addition, the chromosome is laden with specific mobile elements. This includes a prophage located between two tRNA genes (positions 1,347–1,419 kb; length 71 kb) characterized by an entire set of phage structural genes and an integrase gene at the end of the locus, as well as a putative integron (positions 352–394kb; length 41 kb) characterized by an integrase gene at the end of the locus, next to a tRNA gene. The prophage carries no gene with identified function other than phage-related ones, whereas the putative integron carries several genes involved in transport and metabolism.

*'Pseudorhizobium banfieldiae'* (*Pban*)

There are exactly 100 genes exclusively present in *Pban* compared to other *Pseudorhizobium* species (see 'cladeB' in Sup. Table S3). Among them, the main feature is a locus grouping genes encoding the RuBisCO and other functions of the Calvin cycle, as well as respiratory chain cytochromes. These systems allow the provision of electrons and the assimilation of carbon dioxide and are likely to be the main determinants of chemolithoautotrophy in this species. This locus belongs to a larger *Pban*-specific region made of two closely located 27-kb and 80-kb fragments, which suggests it results from the fairly recent insertion and domestication of a mobile element (likely interrupted by a more recent insertion/rearrangement). Among genes in this extended region, several code for enzymes of the classes oxidoreductase, monooxygenase, decarboxylase and glutathione-S transferase, which all use reduced electron acceptors and/or protons; their putative substrates include aromatic cycles and halogenated organic compounds. This suggests a functional link between chemoautotrophy and detoxification pathways. Based on the reconstruction of horizontal gene transfer scenarios, these genes are inferred to originate from a deep-branching lineage of *Neorhizobium* (Sup. Fig. S4). The reconstructed scenarios of HGT (Sup. Table S4) indicate that *Pban* was the recipient of chains of transfer events, dated as early as the diversification of the *Neorhizobium/Pseudorhizobium* group. This suggests genes coding for autotrophy have been circulating for a long time in this group of species, and were only later fixed in the *Pban* lineage.

Other putative functions of *Pban*-specific genes include a biosynthetic locus of a lipopolysaccharide O-antigen with an N-acetylneuramidic acid function. In addition, the genes encoding an imidazole-glycerol-phosphate synthase HisFH present in this locus may operate a modification of the lipopolysaccharide. However, a similar genetic make-up is found in the B-band lipopolysaccharide biosynthesis locus of *Pseudomonas aeruginosa* PAO1, for which the *hisFH* genes have been shown to be dispensable (Rocchetta et al. 1999). These genes show significant similarity to sequences in the phylogenetically distant organisms *Nitrococcus mobilis* Nb-231 (*Gammaproteobacteria*) and *Hoeflea phototrophica* DFL-43 (*Rhizobiales*) (41.8% and 49% average protein identity on 14 and 12 proteins, respectively).

A 26-kb *Pban*-specific region encodes enzymes and transporters related to taurine utilization pathways and to the degradation of possibly halogenated aromatic compounds. These specific functions are reflected by the enrichment of GO terms associated with specific genes with respect to those found in the species' core or pan-genome. These include: 'carbohydrate metabolic process' (GO:0005975) and 'pentose-phosphate shunt' (GO:0006098) that are associated with the RuBisCO operon, and 'N-acetylglucosamine metabolic process' (GO:0006044), 'lipopolysaccharide biosynthetic process' (GO:0009103) and 'histidine biosynthetic process' (GO:0000105) that are associated to the O-antigen biosynthetic locus (Sup. Table S5).

In contrast, there are several genomic regions that have been lost in the *Pban* lineage, including 17 specifically absent genes with respect to cognate *Pseudorhizobium* species. These include an operon present in sister species *Phalo*, *Pfla* and *Pendo* that encodes a multimeric $Na^+/H^+$ cation antiporter (also present in *Pmari* strain MGL06$^T$; gene trees indicate it is a HGT recipient from *Phalo*). Another notable *Pban*-specific gene loss is an operon encoding a cellulose synthase, indicating the likely presence of a cellulose-like polymer in the capsular exopolysaccharide of all other *Pseudorhizobium* species.

Finally, *Pban* genomes specifically lack genes coding for a respiratory complex including several cytochrome c oxidases, in linkage with a gene coding the EutK carboxysome-like microcompartment protein, which known homologues are involved the degradation of ethanolamine. This locus is present in *Pfla, Phalo, Pendo* and *Pmari*, and estimated evolution scenarios for these gene families support a gain by the *Pseudorhizobium* common ancestor (species tree node N9) and a subsequent loss by the *Pban* ancestor (species tree node N15) (Sup. Table S4). Interestingly, the cytochrome c-like enzyme-coding genes included in this locus vary with the strain: *Pfla* YW14 presents a copper-containing nitrite reductase, while *Rhalo* strains AB21 and Khangiran2 present a (non-homologous) secreted nitrous-oxide reductase; *Rendo* strains present no cytochrome c-like genes, but others encoding metabolic enzymes that differ among strains: an amidase, pyridoxamine 5'-phosphate oxidase and alkylhydroperoxidase associated with large metabolite transporters in strain JC140, and carries a cystathionine beta-lyase borne by a transposable element. This suggests that this respiratory chain and associated putative micro-compartment are used as an evolutionary flexible platform for the reductive activities of these organisms.

Note that the nitrite reduction activity is however present in NT-26 (Andres et al. 2013), being coded by a homologous gene that is part of the denitrification pathway locus, shared by all organisms studied here.

*'Pseudorhizobium halotolerans'* (*Phalo*)

This species, represented by strains AB21$^T$ and Khangiran2, features 32 unique genes when compared to all other *Pseudorhizobium* species, including *Pfla*; 86 genes are specific to *Phalo* when compared to *Pfla* and closest species *Pban*; 234 genes are specific to *Phalo* when compared to only sister strain *Pfla* YW14. Enriched GO terms revealed a *Phalo*-specific pathway involved in the biosynthesis of carotenoids (GO:0016117, GO:0006696, GO:0008299; encoded in strain AB21 by genes RHAB21_04118–04126) and biosynthesis of pyridoxine (GO:0008615), a precursor of the vitamin thiamine, as well as signal transduction systems involved in the activation of the former pathways (Sup. Table S5).

*'P. halotolerans+P. flavum'* (*Phalo+Pfla*)

102 genes are specific to the clade and 47 genes are unique to this group within the *Pseudorhizobium* genus (Sup. Table S3, 'clade10' and 'cladeC', respectively). Enriched GO terms revealed *Phalo*-specific pathways linked to the modification of the plasma membrane, such as 'polyketide metabolic process' (GO:0030638), 'cardiolipin biosynthetic process' (GO:0032049) and 'lipoprotein biosynthetic process' (GO:0042158) (Sup. Table S5). These processes are likely to have significant physiological impact, and may play a role in the resistance to high salt concentrations.

'*Pseudorhizobium endolithicum' (*Pendo*)

*Pendo* has apparently undergone a large turnover of its core genome as it diverged from its sister species, with 248 genes unique to this species with respect to the rest of the genus (435 when opposed to the sister clade grouping *Pendo* and *Phalo*). This large set of species-specific gene can be summarized by the enrichment of functions associated with the response to antibiotics (GO:0046677), antibiotic catabolism (GO:0017001), sulphate transport (GO:0008272), urea metabolism (GO:0019627), flagellar motility (GO:0071973) and response to UV light (GO:0009411). In particular, some of the genes coding for flagellar hook-length control protein FliK and flagellin proteins in *Pendo* are highly divergent from those present in other *Pseudorhizobium* species (red sequence labels in flagellin tree, Sup. Fig. S7), and were inferred to be acquired by transfer in the species ancestor. This suggests that this species might exhibit a flagellum-dependent motility different from its relatives within *Pseudorhizobium*, which may have promoted the colonization of rock matrix by *Pendo* (Parag et al. 2013).

Another diagnostic trait of the two available *Pendo* genomes is the clade-specific absence of 299 genes, that are otherwise present in *Pban*, *Phalo*, and *Pfla*. Among these, large contiguous loci are absent from *Pendo*, *with* some segments also being conserved in *Pmari* (i.e. *Pendo* species-specific absence, 132 genes). This pattern indicates that these loci were likely present in the genus ancestor and were lost in the *Pendo* lineage, with some gene homologs being convergently lost in the *Pmari* lineage. Among them, a major feature is the loss of a 30-kb locus that is seemingly involved in the transport and metabolism of aminated carbohydrates, as it includes genes coding for a salicylaldehyde dehydrogenase, a phosphoenolpyruvate synthase and a methylcrotonoyl-CoA carboxylase. A similar pattern is observed for 35 genes interspersed in an 84-kb region (out of 72 genes in this region) that encode various functions including sulphate ion export, carbohydrate metabolism and signal transduction. Finally, *Pendo* specifically lacks a standalone cyanase gene that is otherwise conserved in the genus. The loss of these functions, which are otherwise trademarks of *Pseudorhizobium*, may have followed the ecological niche restriction imposed by the adaptation of *Pendo* to its peculiar lifestyle within rock (Parag et al. 2013).

‘*Pseudorhizobium marinum*’ (*Pmari*)

There are 685 genes specific to *Pmari*, reflecting its basal, highly-diverged position within the *Pseudorhizobium* tree. Among the many functions covered (Sup. Table S5), the following processes are enriched: biosynthesis of polysaccharide (GO:0000271), transport of chloride (GO:0006821), potassium (GO:0006813) and amino acids (GO:0006865), and signal transduction (GO:0007165), notably for regulation of nitrogen utilization (GO:0006808). This notably includes a 27-kb locus containing genes coding for a potassium-transporting ATPase, extrusion transporters and degradation enzymes with putative phenolic compound substrates, and a poly(3-hydroxybutyrate) (PHB) depolymerase.

‘*Pseudorhizobium sub-clade Pban+Phalo+Pfla*’

The largest specific features of the *Pban+Phalo+Pfla* clade are the presence of the 20-kb super-operon *paa* coding for the uptake and degradation of phenylacetate, and of a 13-kb locus including the *sox* operon that encodes the sulphur (thiosulphate) oxidation pathway.

‘*Pseudorhizobium sub-clade Pban+Phalo+Pfla+Pendo*’

Cellular process enriched in the functional annotation of genes specific to the *Pban+Phalo+Pfla+Pendo* clade include NAD cofactor biosynthesis (GO:0009435), tryptophan catabolism (GO:0019441) and phosphatidic acid biosynthesis (GO:0006654) (Sup. Table S5). Other clade-specific genes include an operon coding a thiosulphalte sulphurtransferase with a pyrroloquinoline quinone (PQQ)-binding motif, a SoxYZ-like thiosufalte carrier, a SoxH-like metallo-protease related to beta-lactamases, an inositol monophosphatase and an ABC-type transporter of ferric iron ion. In addition, another clade-specific gene specific in another locus codes for a membrane-bound PQQ-dependent dehydrogenase with glucose, quinate or shikimate as predicted substrates. A 17-kb locus including *pqqBCDE* operon for the biosynthesis of cofactor PQQ and PQQ-dependent methanol metabolism enzymes, was also specifically gained in this clade, but later lost by *Pfla* strain YW14. These genes seem to collectively code for a pathway where the periplasmic oxidation of thiosulphate provides electrons that are carried by SoxYZ and PQQ and used by the metallo-protease and membrane-bound dehydrogenase, respectively, to degrade targeted compounds. The role of the monophosphatase and ferric iron transporter in this process is unclear.

*Pseudorhizobium* (*Pban+Phalo+Pfla+Pendo+Pmari*)

In comparison with the closely related genus *Neorhizobium, Pseudorhizobium*-specific genes are over-represented in genes involved in cellular processes related to energy metabolism: ‘aerobic respiration’ (GO:0009060) and ‘electron transport coupled proton transport’ (GO:0015990), and to anabolic processes, including the biosynthesis of cofactor NAD (GO:0009435), lipid precursor acetyl-CoA from acetate (GO:0019427) and amino-acid asparagine (GO:0006529) (Sup. Table S5).

In addition, a *Pseudorhizobium*-specific operon encodes the biosynthesis of osmoprotectant N-acetylglutaminylglutamine (NAGGN) and an O-antigen ligase domain protein which funtion may be either to further modify NAGGN into a more complex metabolite or to attach it as a decoration of the lipopolysaccharide, even though the latter would likely not provide osmoprotection.

**Verification of bioinformatic predictions of phenotypes**

Predicted clade-specific phenotypic traits for which the experimental test did not match the expectations are reported below.

Pban-*specific use of taurine.* Contrary to the *in silico* prediction, *Pban* strains did not grow in a minimum salts medium with taurine as the sole electron donor under the tested conditions (data not shown). This does not rule out that the strains can use some (possibly other) sulphonate compound, notably under different inducing conditions; further investigation of this trait needs to be undertaken.

Pban+Phalo+Pfla-*specific degradation of phenylacetate*. A screen using the API 20 NE identification system showed *Phalo* and *Pfla* strains were positive for phenylacetate assimilation (Sup. Table S10), in line with *in silico* predictions. However, *Pendo* strain JC140 is also positive, and *Pban* strains are negative, contrary to expectations based on the presence of the *paa* operon. The fact that all positive strains but *Phalo* AB21 had a weak response suggests that induction of this function may not be optimal, or that enzymes encoded by this operon may have low affinity for phenylacetate, questioning the accuracy of their automatic functional annotation. The activity observed in *Pendo* JC140 can be explained by the presence of an isolated gene coding for a 4-hydroxyphenylacetate 3-monooxygenase, the first step in the phenylacetate degradation pathway, which is not homologous to the one present in the *Pban+Phalo+Pfla*-specific *paa* operon. This gene however also occurs in the *Pendo* strain Q54, which is negative in the assimilation test. Based on our limited data, we can only speculate that, in strain Q54 and in *Pban* strains, the identified phenylacetate degradation genes could have lost their function or not be induced in the culture conditions used in the API identification system.

**Metabolic characteristics of the genus *Pseudorhizobium***

Tested on the BioLog GenIII (Sup. Table S8) and API 20 NE systems (Sup. Table S10), strains from member species *P. marinum, P. halotolerans, P. flavum, P. endolithicum and P. banfildiae* were able to utilize D-cellobiose, turanose, N-acetyl-D-glucosamine, D-arabinose, D-glucose, D-mannose, D-fructose, D-galactose, D-mannitol, D-arabitol, myo-inositol, D-glucuronic acid, quinic acid, methyl pyruvate, acetic acid. Strains were not able to utilize stachyose, N-acetyl-b-D-mannosamine, N-acetyl-D-galactosamine, N-acetyl-neuraminic acid, D-raffinose, D-melibiose, inosine, D-glucose-6-phosphate, D-fructose-6-phosphate, D-aspartic acid, D-serine, L-malic acid, bromo-succinic acid. Growth was observed in presence of sodium lactate and rifamycin SV.

#### References


Alexa A, Rahnenfuhrer J. 2010. *topGO: Enrichment Analysis for Gene Ontology*. http://bioconductor.org/packages/topGO/ (Accessed November 30, 2018).

Andres J et al. 2013. Life in an arsenic-containing gold mine: genome and physiology of the autotrophic arsenite-oxidizing bacterium Rhizobium sp. NT-26. Genome Biol Evol. 5:934–953. doi: 10.1093/gbe/evt061.

Bigot T, Daubin V, Lassalle F, Perrière G. 2013. TPMS: a set of utilities for querying collections of gene trees. BMC Bioinformatics. 14:109. doi: 10.1186/1471-2105-14-109.

Camacho C et al. 2009. BLAST+: architecture and applications. BMC Bioinformatics. 10:421. doi: 10.1186/1471-2105-10-421.

Collins C, Didelot X. 2018. A phylogenetic method to perform genome-wide association studies in microbes that accounts for population structure and recombination. PLOS Computational Biology. 14:e1005958. doi: 10.1371/journal.pcbi.1005958.

Doyon J-P, Ranwez V, Daubin V, Berry V. 2011. Models, Algorithms and Programs for Phylogeny Reconciliation. Brief Bioinform. 12:392–400. doi: 10.1093/bib/bbr045.

Fu L, Niu B, Zhu Z, Wu S, Li W. 2012. CD-HIT: accelerated for clustering the next-generation sequencing data. Bioinformatics. 28:3150–3152. doi: 10.1093/bioinformatics/bts565.

Jones P et al. 2014. InterProScan 5: genome-scale protein function classification. Bioinformatics. 30:1236–1240. doi: 10.1093/bioinformatics/btu031.

Lartillot N, Brinkmann H, Philippe H. 2007. Suppression of long-branch attraction artefacts in the animal phylogeny using a site-heterogeneous model. BMC Evolutionary Biology. 7:S4. doi: 10.1186/1471-2148-7-S1-S4.

Lassalle F et al. 2017. Ancestral Genome Estimation Reveals the History of Ecological Diversification in Agrobacterium. Genome Biol Evol. 9:3413–3431. doi: 10.1093/gbe/evx255.

Lassalle F, Jauneikaite E, Veber P, Didelot X. 2019. Automated reconstruction of all gene histories in large bacterial pangenome datasets and search for co-evolved gene modules with Pantagruel. bioRxiv. 586495. doi: 10.1101/586495.

Lassalle F, Muller D, Nesme X. 2015. Ecological speciation in bacteria: reverse ecology approaches reveal the adaptive part of bacterial cladogenesis. Research in Microbiology. 166:729–741. doi: 10.1016/j.resmic.2015.06.008.

Lepage T, Bryant D, Philippe H, Lartillot N. 2007. A General Comparison of Relaxed Molecular Clock Models. Mol Biol Evol. 24:2669–2680. doi: 10.1093/molbev/msm193.

Parag B, Sasikala C, Ramana CV. 2013. Molecular and culture dependent characterization of endolithic bacteria in two beach sand samples and description of Rhizobium endolithicum sp. nov. Antonie Van Leeuwenhoek. 104:1235–1244. doi: 10.1007/s10482-013-0046-7.

Rocchetta HL, Burrows LL, Lam JS. 1999. Genetics of O-Antigen Biosynthesis inPseudomonas aeruginosa. Microbiol. Mol. Biol. Rev. 63:523–553.

Ronquist F, Huelsenbeck JP. 2003. MrBayes 3: Bayesian phylogenetic inference under mixed models. Bioinformatics. 19:1572–1574.

Seemann T. 2014. Prokka: rapid prokaryotic genome annotation. Bioinformatics. 30:2068–2069. doi: 10.1093/bioinformatics/btu153.

Shams M, Vial L, Chapulliot D, Nesme X, Lavire C. 2013. Rapid and accurate species and genomic species identification and exhaustive population diversity assessment of Agrobacterium spp. using recA-based PCR. Syst. Appl. Microbiol. 36:351–358. doi: 10.1016/j.syapm.2013.03.002.

Sievers F et al. 2011. Fast, scalable generation of high-quality protein multiple sequence alignments using Clustal Omega. Molecular Systems Biology. 7:539. doi: 10.1038/msb.2011.75.

Stackebrandt E et al. 2002. Report of the ad hoc committee for the re-evaluation of the species definition in bacteriology. Int. J. Syst. Evol. Microbiol. 52:1043–1047.

Stamatakis A. 2014. RAxML version 8: a tool for phylogenetic analysis and post-analysis of large phylogenies. Bioinformatics. btu033. doi: 10.1093/bioinformatics/btu033.

Steinegger M, Söding J. 2017. MMseqs2 enables sensitive protein sequence searching for the analysis of massive data sets. Nature Biotechnology. doi: 10.1038/nbt.3988.

Suyama M, Torrents D, Bork P. 2006. PAL2NAL: robust conversion of protein sequence alignments into the corresponding codon alignments. Nucleic Acids Research. 34:W609–W612. doi: 10.1093/nar/gkl315.

Suzuki R, Shimodaira H. 2006. Pvclust: an R package for assessing the uncertainty in hierarchical clustering. Bioinformatics. 22:1540–1542. doi: 10.1093/bioinformatics/btl117.

Szöllősi GJ, Rosikiewicz W, Boussau B, Tannier E, Daubin V. 2013. Efficient Exploration of the Space of Reconciled Gene Trees. Syst Biol. 62:901–912. doi: 10.1093/sysbio/syt054.

Szöllősi GJ, Tannier E, Daubin V, Boussau B. 2015. The inference of gene trees with species trees. Syst. Biol. 64:e42–e62. doi: 10.1093/sysbio/syu048.

Szöllősi GJ, Tannier E, Lartillot N, Daubin V. 2013. Lateral Gene Transfer from the Dead. Syst Biol. 62:386–397. doi: 10.1093/sysbio/syt003.
