## Supplementary material for "Phylogenomic analysis reveals the basis of adaptation of *Pseudorhizobium* species to extreme environments": Table 3

| | |
|---|---|
| **TAXONUMBER** | TA00814 |
| **TYPE OF DESCRIPTION** | New Description |
| **SPECIES NAME** | Pseudorhizobium banfieldii |
| **GENUS NAME** | Pseudorhizobium |
| **SPECIFIC EPITHET** | banfieldii |
| **SPECIES STATUS** | sp. nov. |
| **SPECIES ETYMOLOGY** | in honour of Prof Jill Banfield, environmental microbiologist whose research revolutionized the view of bacterial and archeal diversity. |
| **AUTHORS** | Lassalle F, Osborne T, Brinkmann H, Dastgheib M, Zhao F, Zhang J, Balloux F, Petersen J, Santini JM |
| **SUBMITTER** | FLORENT LASSALLE |
| **E-MAIL OF THE SUBMITTER** | |
| **DESIGNATION OF THE TYPE STRAIN** | NT-26 |
| **STRAIN COLLECTION NUMBERS** | DSM 106348 |
| **16S rRNA GENE ACCESSION NUMBER** | AF159453 |
| **GENOME ACCESSION NUMBER [RefSeq]** | GCF_000967425.1 |
| **GENOME STATUS** | complete |
| **GENOME SIZE** | 4577425 |
| **GC mol %** | 61.84 |
| **COUNTRY OF ORIGIN** | Australia |
| **REGION OF ORIGIN** | Northern Territory |
| **DATE OF ISOLATION UNKNOWN (< yyyy)** | < 1999 |
| **SOURCE OF ISOLATION** | moist arsenopyrite-containing rock |
| **SAMPLING DATE** | 1999-01-01 |
| **GEOGRAPHIC LOCATION** | Granites Gold Mine |
| **LATITUDE** | 20°32′18.4″S |
| **LONGITUDE** | 130°18′37.7″E |
| **DEPTH** | 60 |
| **NUMBER OF STRAINS IN STUDY** | 3 |
| **SOURCE OF ISOLATION OF NON-TYPE STRAINS** | moist arsenopyrite-containing rock, soil |
| **GROWTH MEDIUM, INCUBATION CONDITIONS [Temperature, pH, and further information] USED FOR STANDARD CULTIVATION** | Minimal salts medium (MSM) as per (Santini et al., 2000. Applied & Environmental Microbiology 66(1):92-97).<br>pH 8.0<br>28°C |
| **IS A DEFINED MEDIUM AVAILABLE** | yes (Santini et al., 2000. Applied & Environmental Microbiology 66(1):92-97). |
| **ALTERNATIVE MEDIUM 1** | Luria Bertani |
| **GRAM STAIN** | NEGATIVE |
| **CELL SHAPE** | rod |
| **CELL SIZE (length or diameter)** | 1.0 |
| **MOTILITY** | motile |
| **IF MOTILE** | flagellar |
| **IF FLAGELLATED** | 2 sub-terminal flagella |
| **SPORULATION (resting cells)** | none |
| **COLONY MORPHOLOGY** | produces EPS |
| **LOWEST pH FOR GROWTH** | 4 in MSM + arsenite |
| **HIGHEST pH FOR GROWTH** | 9 in MSM + arsenite |
| **pH OPTIMUM** | 8.0 in MSM + arsenite |
| **pH CATEGORY** | neutrophile |
| **RELATIONSHIP TO O2** | facultative aerobe |
| **O2 CONDITIONS FOR STRAIN TESTING** | aerobiosis |
| **CARBON SOURCE USED [class of compounds]** | sugars, organic acids, carbon dioxide |
| **CARBON SOURCE USED [specific compounds]** | acetate, arabinose, galactose, fructose, fumarate, glucose, glycerol, inositol, lactate, lactose, malate, maltose, mannitol, pyruvate, trehalose, raffinose, salicin, succinate, sucrose, xylose |
| **CARBON SOURCE NOT USED [specific compounds]** | citrate, rhamnose, sorbitol. |
| **NITROGEN SOURCE** | NO3, NH4+ |
| **TERMINAL ELECTRON ACCEPTOR** | oxygen, nitrate, nitrite |
| **ENERGY METABOLISM** | mixotroph |
| **BIOSAFETY LEVEL** | 1 |
| **HABITAT** | ENVO:00001995, ENVO:00001998, ENVO:00005801 |
| **BIOTIC RELATIONSHIP** | free-living |
| **KNOWN PATHOGENICITY** | none |
| **MISCELLANEOUS, EXTRAORDINARY FEATURES RELEVANT FOR THE DESCRIPTION** | facultative chemolithoautotrophe on thiosulfate as electron source and carbon dioxide as C source |

DRAFT DRAFT
